# Spatial Insulation Confines Dynamic Chromosomes in Minimal Marine Eukaryotes

**DOI:** 10.1101/2025.04.21.649811

**Authors:** Martha Valiadi, Keith Harrison, Yann Loe-Mie, Bryony Williams, Dyan Ankrett, Nicholas Smirnoff, Adam Monier

## Abstract

Marine picoeukaryotes of the order Mamiellales, including *Ostreococcus tauri*, the smallest known free-living eukaryote, possess compact genomes yet maintain enigmatic “outlier chromosomes” characterised by lower GC content and hypervariability. To determine the structural and functional nature of these regions, we applied chromosome conformation capture to *O. tauri* and conducted comparative multi-omics analyses across the Mamiellales order, presenting the first analysis of three-dimensional genome organisation in marine picoeukaryotes. We reveal that outlier regions form structures resembling topologically associating domains, with sharp boundaries that spatially insulate them from the standard chromosomes. These compartments are defined by a distinctive chromatin state characterised by hypomethylation and transcriptional hyperactivity, and are frequently, though not universally, enriched in transposable elements. Crucially, species that lack transposable element enrichment in their outlier chromosomes nonetheless retain the transcriptional hyperactivity and distinct nucleotide composition of these regions, indicating that the functional identity of these compartments persists independently of transposon accumulation. The dynamic nature of these insulated domains is highlighted by the presence of structurally diverse giant polyketide synthase loci. We identify convergent genomic organisation in other chlorophytes, as well as phylogenetically distant stramenopiles. Our results suggest that such compartmentalisation of rapidly evolving, dynamic genomic regions represents a fundamental architectural principle of minimal eukaryotic genomes.

## Introduction

The marine green alga *Ostreococcus tauri* is the smallest known free-living eukaryote (∼0.8 μm), containing only a single plastid and mitochondrion (1, 2). This minimal cellular organisation presents a simplified model for studying fundamental eukaryotic processes such as photosynthesis, circadian rhythms, and environmental responses (3–5). *Ostreococcus* is a member of the Mamiellales (class Mamiellophyceae), a group that also includes other picoalgae such as *Bathycoccus*, *Mantoniella*, and *Micromonas* (6). Mamiellales are globally distributed, thriving in diverse marine environments from tropical to polar waters (7–9), supported by ecotype differentiation and intra-order genetic diversity (10–12).

A distinctive feature of Mamiellales genomes is the presence of two “outlier chromosomes”, known as the big and small outlier chromosomes (BOC and SOC), each containing regions of markedly lower GC content (referred to as BOC1 and SOC1) (13). These regions are enriched in introns, transposable elements (TEs), highly expressed genes (14–16), and they vary significantly among *Ostreococcus* strains, with polymorphisms associated with phenotypic differences in growth and viral susceptibility (17, 18). In *O. tauri*, the BOC contains a putative mating-type locus and genes linked to chromatin conformation (14, 17, 19), while the SOC contributes to antiviral defence (18, 20). Although outlier chromosomes have been recognised since the first Mamiellales genomes were sequenced (21), fundamental questions about these unusual genomic regions remain unanswered. How are outlier chromosomes organised in three-dimensional nuclear space? What regulatory mechanisms distinguish them from the rest of the genome? And are their distinctive features conserved across the Mamiellales lineage, or specific to individual species?

We addressed these questions using a multi-omics approach–including chromosome conformation capture (Hi-C), transcriptomics, genome-wide methylation analysis, and metabolomics–to characterise the structural, epigenetic, and functional properties of outlier chromosomes across multiple Mamiellales species. Our DNA interaction analysis, the first in picoeukaryotes, reveals that BOC1 and SOC1 form spatially insulated compartments akin to topologically associating domains (TADs). These regions share conserved features across species: reduced DNA methylation, elevated gene expression, and enrichment of lineage-specific genes including giant type I polyketide synthases. Yet the underlying sequence composition varies markedly: TEs dominate in *Ostreococcus tauri* and *Micromonas* spp. but are largely absent from *O. lucimarinus* and *Bathycoccus prasinos*, which nonetheless retain the distinctive sequence and transcriptional profiles of these compartments. Strikingly, we find similar spatial insulation of equivalent outlier regions in the phylogenetically distant stramenopile *Pelagomonas calceolata*, suggesting this chromosomal organisation has arisen independently in unrelated lineages. Our findings reveal that conserved genomic compartmentalisation, rather than any single sequence-level mechanism, defines these unusual regions in minimal eukaryotic genomes.

## Results

### Outlier regions are conserved across Mamiellales and enriched in genes associated with genomic rearrangement

To determine whether BOC1 and SOC1 represent a conserved feature of Mamiellales genomes, we examined six species with high-quality genome assemblies. Despite their highly compact genomes (SI Appendix, Table S1 and S2), all species examined (*Ostreococcus* sp. RCC809, *O. tauri*, *O. lucimarinus*, *Micromonas commoda*, *M. pusilla*, and *Bathycoccus prasinos*) contain distinct low-GC outlier regions (14, 22–24). Even in *Mantoniella tinhauana*, where their presence had been uncertain (25), we identified two such regions with ∼15% lower GC content than the surrounding genome (SI Appendix, Fig. S1 and Table S2). Although BOC1 and SOC1 regions vary considerably in size across lineages (Fig. 1a), their consistent presence suggests they are an ancient feature of Mamiellales. Comparative analysis revealed that BOC1 regions maintain greater evolutionary conservation than SOC1. While SOC1 regions showed limited synteny and fewer orthologues across genera (SI Appendix, Fig. S2), BOC1 regions shared more orthologues, though not strict synteny (Fig. 1a).

**Figure 1.**
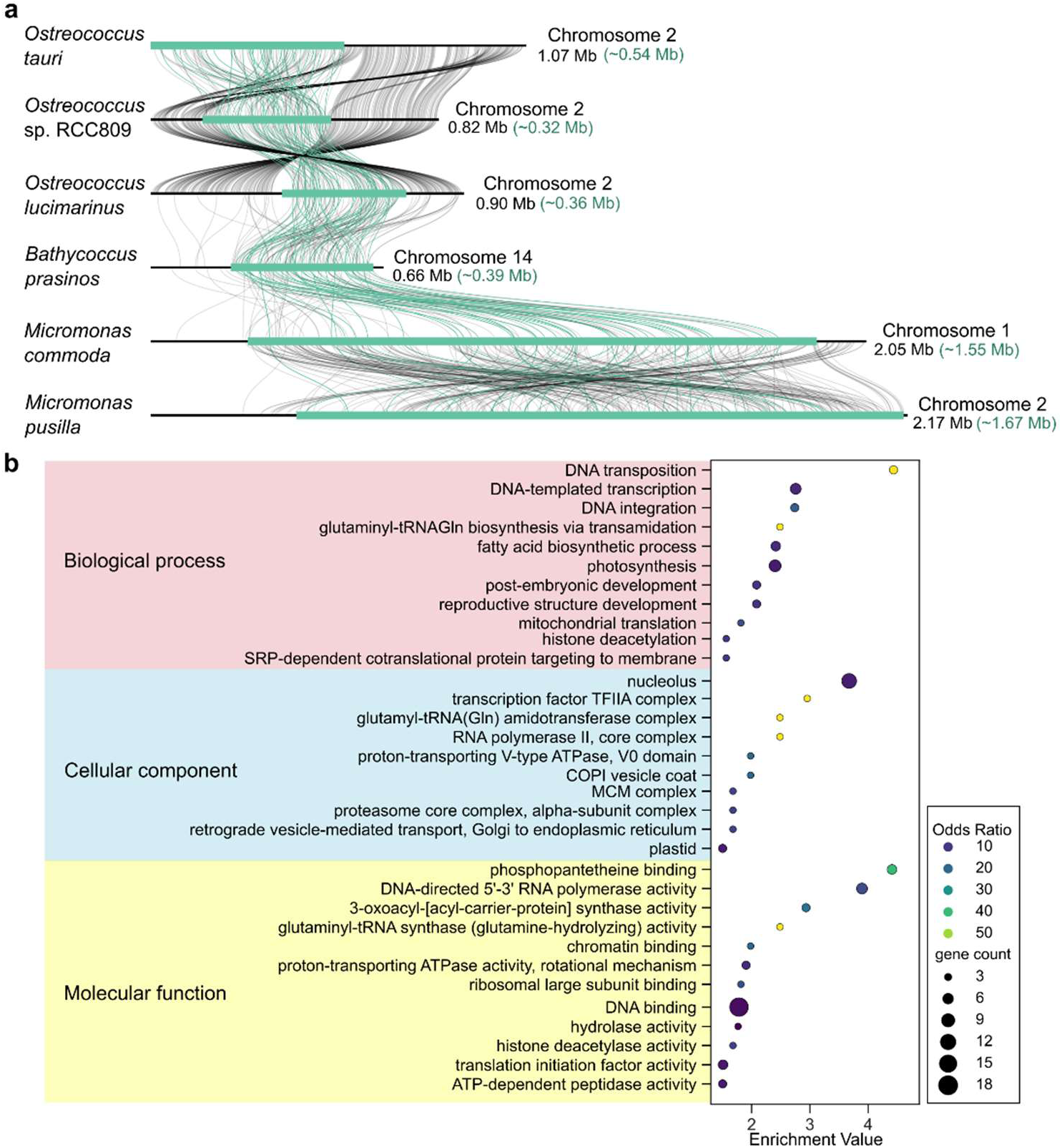
Comparative genomic analysis of BOC1 regions across Mamiellales species. (**a**), Syntenic analysis and gene homology visualisation of BOC1 regions across six Mamiellales species. Connecting lines indicate homologous genes identified through reciprocal best BLASTP hits between *O. tauri* RCC4221, *O.* sp. RCC809, *O. lucimarinus* CCE9901, *B. prasinos* RCC1105, *M. pusilla* CCMP1545, and *M. commoda* RCC299. BOC1 regions are displayed in green. BOC1-localised orthologues are indicated by green lines. Total BOC1 length and the proportion contained within the region (in parentheses) are indicated for each species. (**b**), Gene Ontology (GO) enrichment analysis of BOC1-encoded proteins in RCC4221. The plot displays GO terms categorised into three major classes: biological process (pink), cellular component (blue), and molecular function (yellow). Enrichment values are shown as −Log_10_(p-value), with a significance threshold of p<0.0335 (Fisher’s exact test). Dot size indicates gene count, while colour intensity represents the odds ratio, defined as the proportion of GO term occurrence in BOC1 proteins relative to non-BOC1 regions.

The gene content of these regions suggests a distinct functional profile. Gene Ontology enrichment analysis of *O. tauri* BOC1 revealed a high concentration of genes involved in DNA transposition, transcription, and integration (Fig. 1b; Dataset S1). The region also contains genes for chromatin remodelling, such as mini-chromosome maintenance (MCM) complex components and sister chromatid cohesion protein 2 (SCC2). In contrast, SOC1 is enriched for genes related to RNA polymerase activity, membrane components, and methylation machinery (Dataset S2), indicating distinct functional profiles between the two outlier regions.

### Transposable elements and introns cluster in outlier genomic regions across Mamiellales genomes

We conducted a survey of introns and TEs across Mamiellales genomes, confirming previous findings (21, 24–26) and revealing a distinct, non-random distribution of these features. In *O. tauri*, BOC1 and SOC1 contain 14.5-fold and 17.4-fold more TEs, respectively, than the standard genome (Kolmogorov–Smirnov test, p<0.001; Fig. 2a,b; SI Appendix, Fig. S3; Dataset S3). For example, Ty1/Copia retrotransposon density increases from 2.04 elements per Mb in the standard genome to 37.4 and 167 elements per Mb in BOC1 and SOC1, respectively. Similar patterns are observed across most Mamiellales species (Dataset S4).

**Figure 2.**
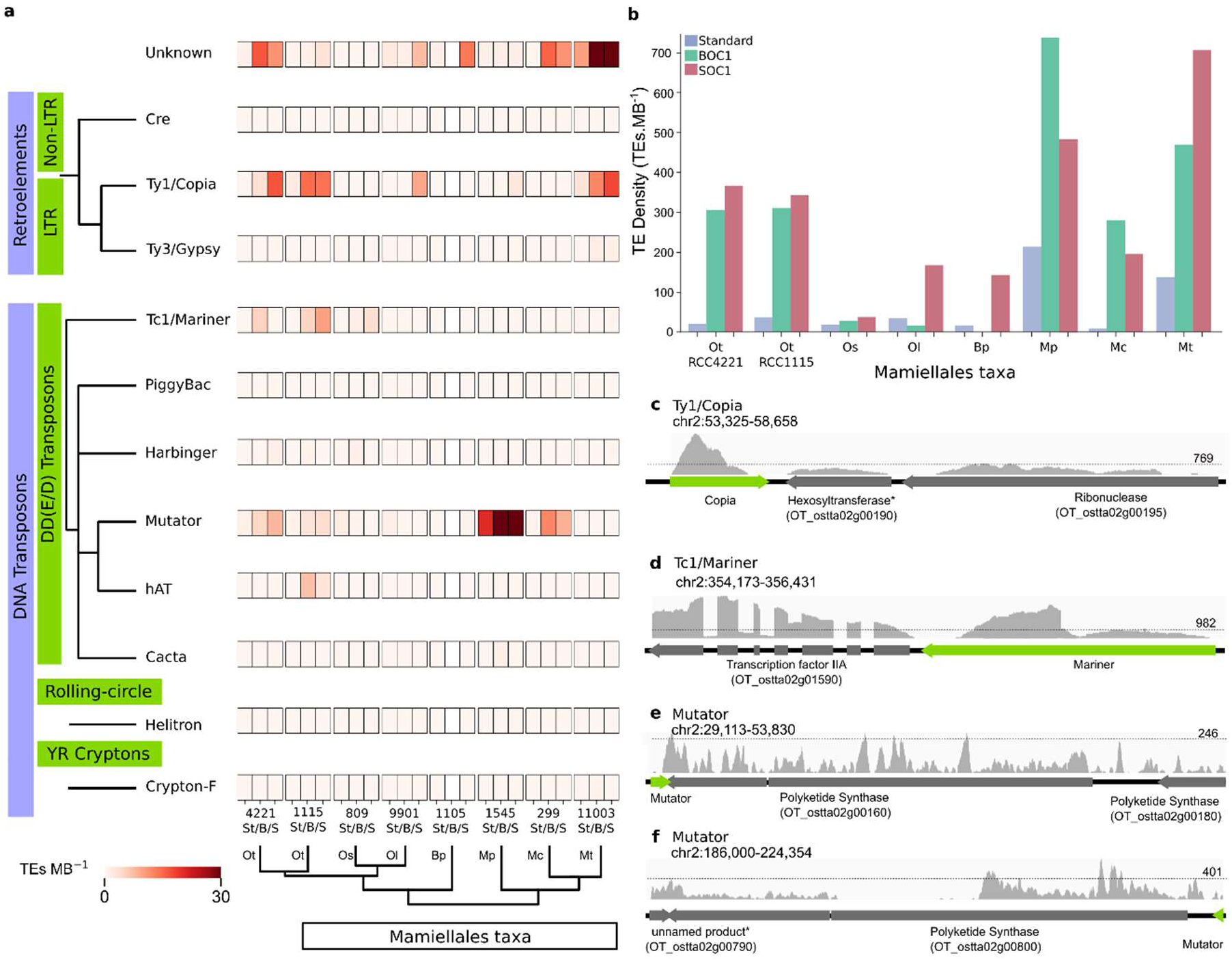
Distribution and expression of transposable elements across Mamiellales genomes. (**a**), Dendrogram of major transposable element (TE) classes present in Mamiellales. The left sidebars indicate the classification of elements into retrotransposon, with further subdivision into LTR/non-LTR retrotransposons and DDE/D transposons, rolling-circle elements, and YR cryptons. Heatmap shows TE density (TEs Mb⁻¹) across standard (St), BOC1 (B), and SOC1 (S) genomic regions. The lower dendrogram displays phylogenetic relationships between Mamiellales species: *O. tauri* (Ot; strains RCC4221 and RCC1115), *O.* sp. RCC809 (Os), *O. lucimarinus* (Ol; CCE9901), *B. prasinos* (Bp; RCC1105), *M. pusilla* (Mp; CCMP1545), *M. commoda* (Mc; RCC299), and *M. tinhauana* (Mt; RCC11003). (**b**), Quantitative comparison of TE abundance (TEs Mb⁻¹) across different genomic regions. Bar colours correspond to standard (grey), BOC1 (green), and SOC1 (red) regions. (**c—f**), RNA-Seq expression profiles of representative TEs in *O. tauri* RCC4221 showing: (c) Ty1/Copia element localised upstream of a putative hexosyltransferase gene; (d) Tc1/Mariner element near a transcription factor IIA; (e and f) Mutator elements near polyketide synthases. Grey peaks represent RNA-Seq coverage, with maximum coverage values indicated in the top right corner of each track. Dotted lines represent 200x RNA-Seq coverage threshold. Grey bars indicate predicted protein-coding regions with their corresponding NCBI locus tags (genome assembly GCF_000214015.3).

We next examined the genomic context of TEs, as they often neighbour actively transcribed genes. In *O. tauri*, transcriptome analysis revealed associations between TEs and transcriptionally active genes in BOC1 and SOC1 (Fig. 2c–f; Dataset S5). For example, a ∼5 kb Ty1/Copia retrotransposon sits adjacent to a hexosyltransferase gene (Fig. 2c), while a ∼1 kb Tc1/Mariner DNA transposon is near a transcription factor IIA gene (Fig. 2d). Mutator-like elements (MULEs) frequently associate with giant modular polyketide synthase (PKS) genes, sometimes inserting upstream and downstream of coding segments (Fig. 2e,f).

Intron distribution shows a similar pattern: BOC1 regions consistently show higher intron density across taxa, for instance four-fold higher than the standard genome in *O. tauri* (Dataset S4).

### Genes on outlier regions are more highly expressed and frequently spliced

Analysis of transcriptomic data from *O. tauri* and other Mamiellales species revealed that genes in BOC1 regions consistently show higher expression compared to those in the standard genome. In *O. tauri*, outlier genes show 3.8-fold higher mean expression (Kruskal-Wallis, p<0.001; Fig. 3a, d). Similar trends are observed in *M. commoda*, *M. pusilla*, and *B. prasinos* (2.4, 3.3, and 1.6-fold, respectively; p<0.001; Fig. 3a–f), although not all species display higher expression in SOC1 (Dataset S6). In most species, this elevated transcription is negatively correlated with GC content, such that lower GC content in outlier regions is associated with higher gene expression (Spearman’s rank, ρ=−0.37, −0.29, and −0.49 for *O. tauri*, *M. commoda*, and *M. pusilla*, respectively; p<0.001; Fig. 3g–i; SI Appendix, Table S3). However, in *B. prasinos*, the trend is reversed, with higher GC content correlating with increased gene expression (ρ=0.22, p<0.001; Fig. 3i).

**Figure 3.**
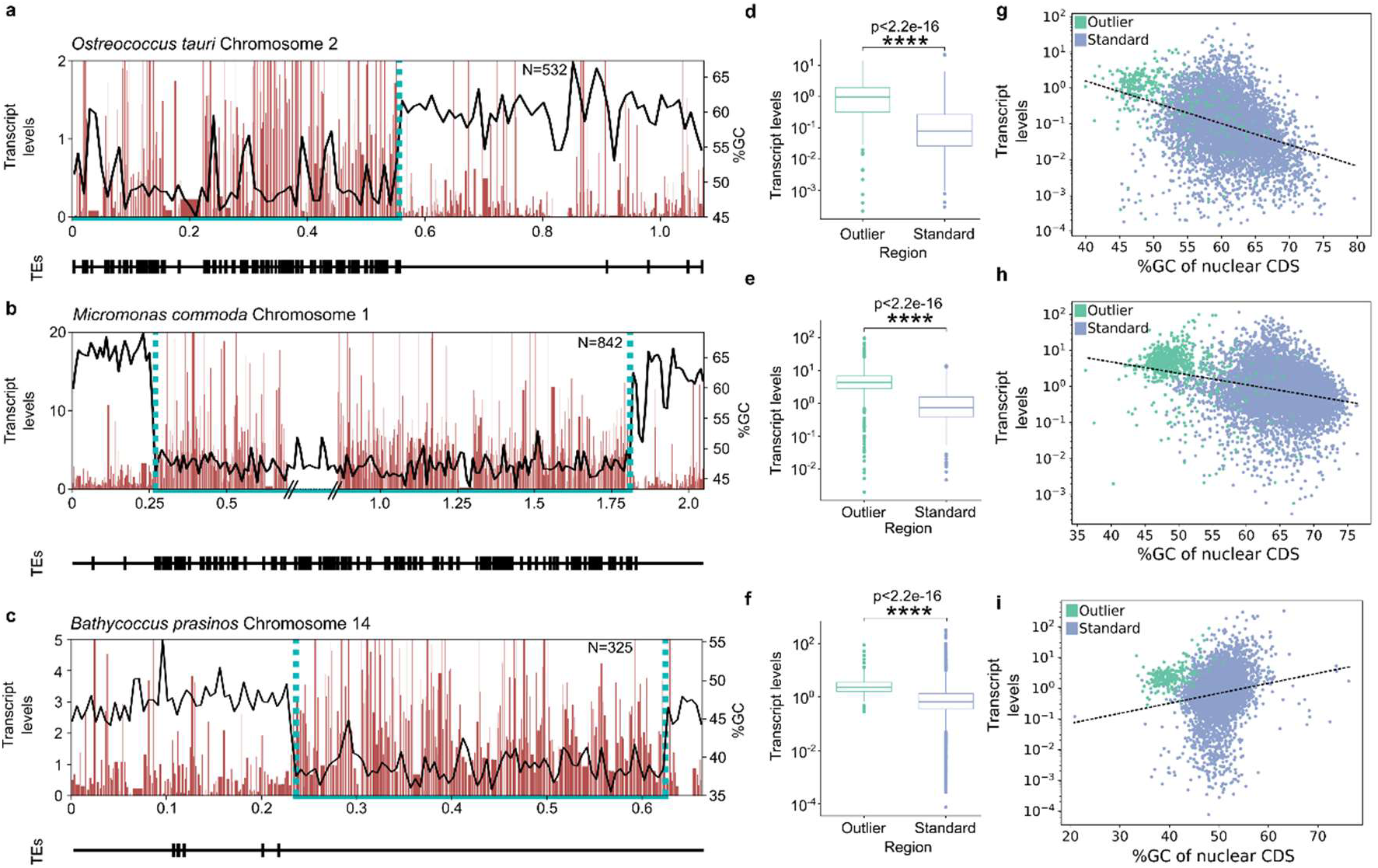
Higher gene expression in BOC1. (**a—c**), Expression levels, GC content and TE content across BOC of (**a**) *O. tauri* (RCC4221; chromosome 2), (**b**), *M. commoda* (RCC299; chromosome 1), and (**c**) *B. prasinos* (RCC1105; chromosome 14). Boundary points of the BOC1 regions are labelled with cyan dotted lines; transposable elements are shown on the lower track. (**d—f**), Comparative analysis of gene expression in outlier regions and standard genome; box plots display the distribution of transcript levels across outlier (BOC1 and SOC1) and standard regions for each taxon (Kruskal-Wallis, p<0.001). (**g—i**), Correlation between gene expression and GC content of coding sequences (CDS) in (**g**), *O. tauri* (Spearman’s rank, ρ=−0.366, p=7.82e^-242^), (**h**), *M. commoda* (ρ = −0.285, p = 3.70e^-192^), and (**i**) *B. prasinos* (ρ = -0.38302, p = 8.49e^-13^); CDS in BOC1 and non-BOC1 regions are represented in green and blue, respectively.

In addition to elevated expression, outlier regions contribute disproportionately to transcriptional diversity through alternative splicing. Although the BOC1 region in *O. tauri* contains far fewer genes than the standard genome, it generates 38% (1,216 of 3,215) of all detected splice variants. Similar enrichment of splice variants is observed in *O. lucimarinus*, *B. prasinos*, and *M. commoda* (20%; Dataset S7).

### Hi-C analysis reveals spatial insulation of outlier regions

To examine the three-dimensional organisation of outlier regions, we performed Hi-C on *O. tauri*. Contact maps (330 million contacts at 10 kb resolution; Fig. 4a; SI Appendix, Fig. S4) showed a simple architecture dominated by intrachromosomal interactions (77% of all contacts; Dataset S8). BOC and SOC displayed disproportionately high intrachromosomal contacts (∼20% of total; Fig. 4; SI Appendix, Fig. S5).

**Figure 4.**
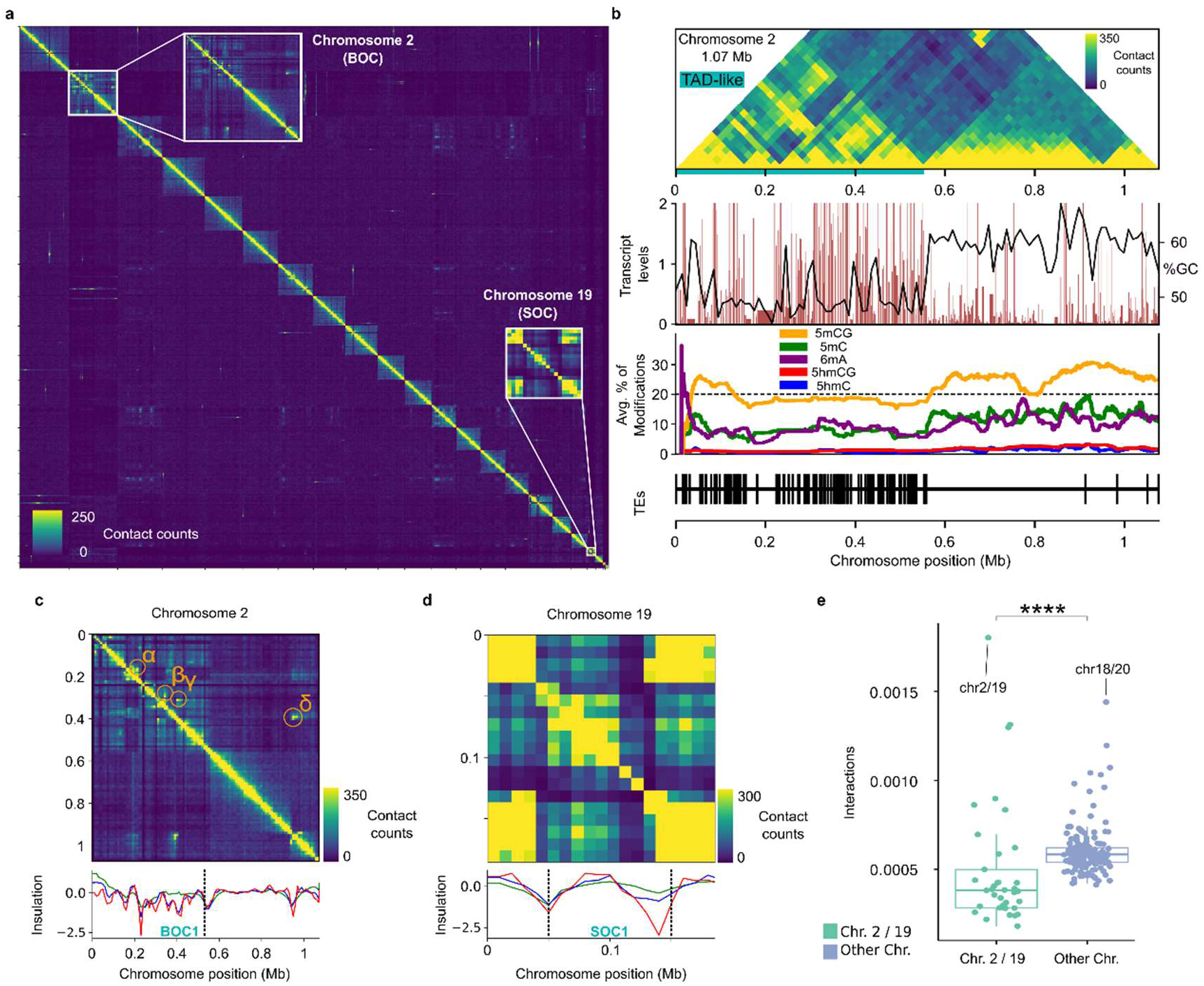
Chromosomal organisation and three-dimensional genome architecture in *O. tauri*. (**a**), Hi-C contact map of the *O. tauri* genome binned at 10 kb resolution with normalised interactions counts; chromosomes 2 and 19, which display enriched intrachromosomal contacts, highlighted by white squares. (**b**), Hi-C normalised contact map of chromosome 2 at 20 kb resolution. The first track displays transcript levels alongside GC content. The second track shows the rolling average of DNA modifications within a scaled window, 5mCG (orange), 5mC (green), 6mA (purple), 5hmCG (blue). The third track indicates positions of transposable elements (black). (**c—d**), Hi-C contact maps binned at 10 kb for (**c**) chromosome 2 and (**d**) chromosome 19. Topologically associating domains are indicated by black dotted lines, with outlier regions labelled as BOC1 and SOC1 (cyan text), the bottom track represents the insulation score calculated as a rolling z-score from the summation of Hi-C contacts at 5 kb (red), 10 kb (blue), and 20 kb (green) resolutions. (**e**), Distribution of normalised interchromosomal contact probabilities between different chromosomes. Interactions involving BOC or SOC chromosomes are labelled in green, whilst interactions between standard nuclear are shown in blue. Notable interactions between chromosomes 2 and 19, and between chromosomes 18 and 20 are highlighted.

BOC1 forms a clear topologically associating domain (TAD)-like structure (Fig. 4b, c; SI Appendix, Fig. S6), bounded by sharp domain borders and containing four putative chromatin loops (α, β, γ, δ). Genes within these loops encode diverse functions, including glycolipid biosynthesis and chromatin binding (Dataset S9). SOC1 exhibits a similar but smaller insulated pattern (Fig. 4d). The Hi-C map of SOC suggests a TAD-like structure encompassing SOC1, although the small size of SOC1 limits the ability to computationally identify domains.

Quantitative analysis showed that BOC and SOC have significantly fewer interchromosomal contacts than standard regions (Fig. 4e; p<0.0001; Dataset S10). Instead, these outlier regions preferentially interact with each other, indicating physical separation from the rest of the genome. Analysis of *M. tinhauana* Hi-C data (25) revealed similar compartmentalisation patterns (SI Appendix, Fig. S1; Dataset S8), suggesting that spatial insulation of outlier regions is conserved across the Mamiellales lineage.

### Outlier regions maintain distinctive epigenetic landscapes

Whole-genome methylation profiling revealed that epigenetic and spatial boundaries are precisely aligned. In *O. tauri*, a sharp shift in methylation at ∼0.55 Mb on chromosome 2 precisely demarcates the BOC1 region (Fig. 4b), aligning with the TAD-like boundary identified by Hi-C and the zone of higher TE density and gene expression. Methylation levels within outlier regions are consistently lower than in the standard genome. BOC1 shows a 1.4-fold reduction in CpG 5-methylcytosine (5mCG) relative to the rest of the genome (Fig. 4b; SI Appendix, Fig. S7 and S8; Dataset S11). Overall, 5-methylcytosine (5mC) and N6-methyladenine (6mA) are reduced by 1.7- and 1.4-fold, respectively, while 5-hydroxymethylcytosine (5hmC) and 5hmCG remain uniformly low across the chromosome. SOC1 displays similarly distinctive patterns, with a sharp rise in 6mA at its start and increased 5mCG at its end (SI Appendix, Fig. S9).

Available bisulfite sequencing data (27) confirmed this pattern across species: *O. lucimarinus* shows CpG methylation declining from 22.2% in standard regions to 19.2% in BOC1 and 16.2% in SOC1. *M. pusilla* shows a similar trend with levels decreasing from 17.8% to 14.5% in BOC1 and 10.9% in SOC1 (Fig. 5a,b; SI Appendix, Fig. S10; Dataset S12).

**Figure 5.**
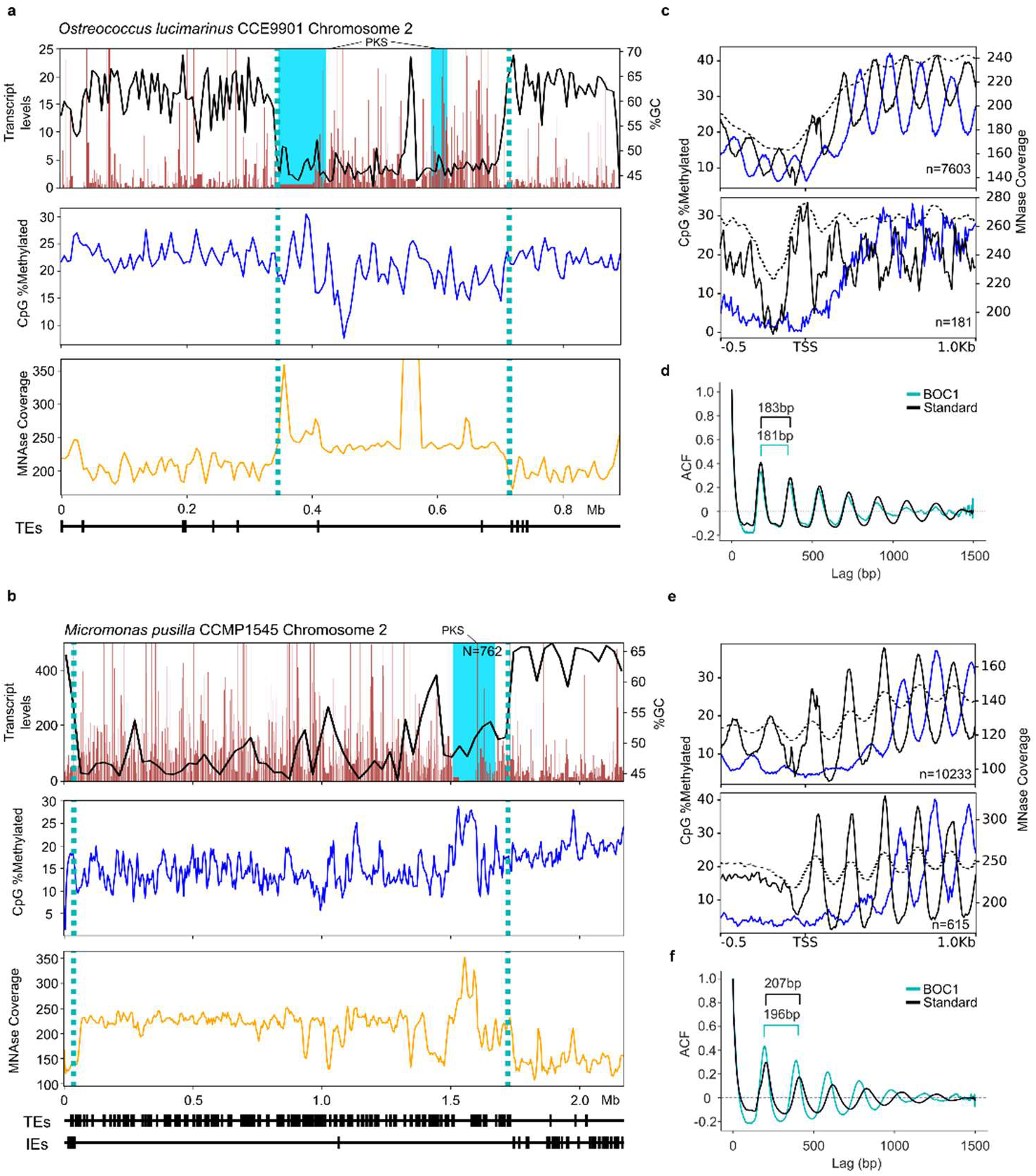
MNase and CpG profiles across BOC1 in *Ostreococcus lucimarinus* CCE9901 and *Micromonas pusilla* CCMP1545. (**a, b**), Transcript levels, CpG methylation as a percentage of base coverage which is methylated, MNase coverage alongside TEs and IEs in *O. lucimarinus* CCE9901 (**a**) and *M. pusilla* CCMP1545 (**b**). Dotted lines indicate the edge of outlier regions. Blue bars indicate PKS coding regions. (**c, e**), Means at each position aligned to transcription start sites are shown for CpG methylation and nucleosomes within the standard genome above and BOC1 genes below for *O. lucimarinus* (**c**) and *M. pusilla* (**e**). Blue is the percentage of bases which are CpG methylated. Black is nucleosome centre counts. Light grey is MNase coverage. (**d, f**), The autocorrelation function estimates of nucleosome centre counts are shown for each lag (offset, bp) across standard genome (black) and BOC1 (turquoise) genes for *O. lucimarinus* (**d**) and *M. pusilla* (**f**), respectively. Period is annotated in base pairs.

### TE density correlates with altered nucleosome spacing and chromatin flexibility

The variation in TE abundance between Mamiellales species provides an opportunity to examine whether TE load correlates with chromatin structure. We re-analysed published MNase-seq data from *O. lucimarinus* and *M. pusilla* (27) to determine how variation in TE density corresponds to nucleosome spacing and positioning within BOC1. Across both species, nucleosome organisation in BOC1 bears hallmarks of transcriptionally permissive chromatin: high nucleosome occupancy around transcription start sites, weaker periodicity in promoters, and reduced CpG methylation (Fig. 5c,e). These features coincide with elevated transcription: RNA-Seq coverage in BOC1 is 1.86× and 2.44× higher than in standard regions for *O. lucimarinus* and *M. pusilla*, respectively (Kruskal–Wallis, p < 0.001).

Nucleosome spacing differs markedly between species and genomic compartments. In the TE-rich *M. pusilla*, nucleosome repeat length shortens from 207 bp in standard regions to 196 bp in BOC1, whereas the TE-poor *O. lucimarinus* shows only a minor shift from 183 bp to 181 bp (Fig. 5d,f). In both species, outlier regions show reduced chromatin stiffness and more well-positioned nucleosomes. Comparisons with invariant chromosomes further support these findings (SI Appendix, Fig. S11; Dataset S13).

TE–nucleosome spatial relationships further distinguish these regions. Whereas *O. lucimarinus* BOC1 shows little TE enrichment, *M. pusilla* BOC1 contains 3.5× more TEs than its standard genome. Its BOC1 also shows a depletion in introner elements (IEs), which are otherwise abundant outside this region (24, 28). TEs display a bimodal distribution with peaks ±50 bp from nucleosome centres, especially pronounced in BOC1, consistent with preferential insertion into accessible linker DNA rather than nucleosome cores (SI Appendix, Fig. S12). Notably, the same regions encoding giant type I PKS genes exhibit elevated MNase coverage and CpG methylation (Fig. 5a,b).

### Giant type I polyketide synthase genes show structural diversity with distinct metabolic outputs

Giant type I PKS genes, a conserved feature of BOC1 regions, show large structural variations across Mamiellales species. PKS gene architecture varies dramatically, with lengths ranging from 14,393 bp in *O. tauri* to 64,971 bp in *M. tinhauana* (Fig. 6a; Dataset S14). Even within species, intraspecific variation is substantial: different *O. tauri* strains harbour PKS genes ranging from 14,393 to 33,947 bp. Long-read resequencing of our *O. tauri* RCC4221 laboratory culture (“Exeter”) revealed that the overall BOC1 architecture remains conserved relative to the reference genome, but the PKS locus displays substantial structural reorganisation, with additional modules, domain rearrangements, and multiple inversions (Fig. 6a; Dataset S14).

**Figure 6.**
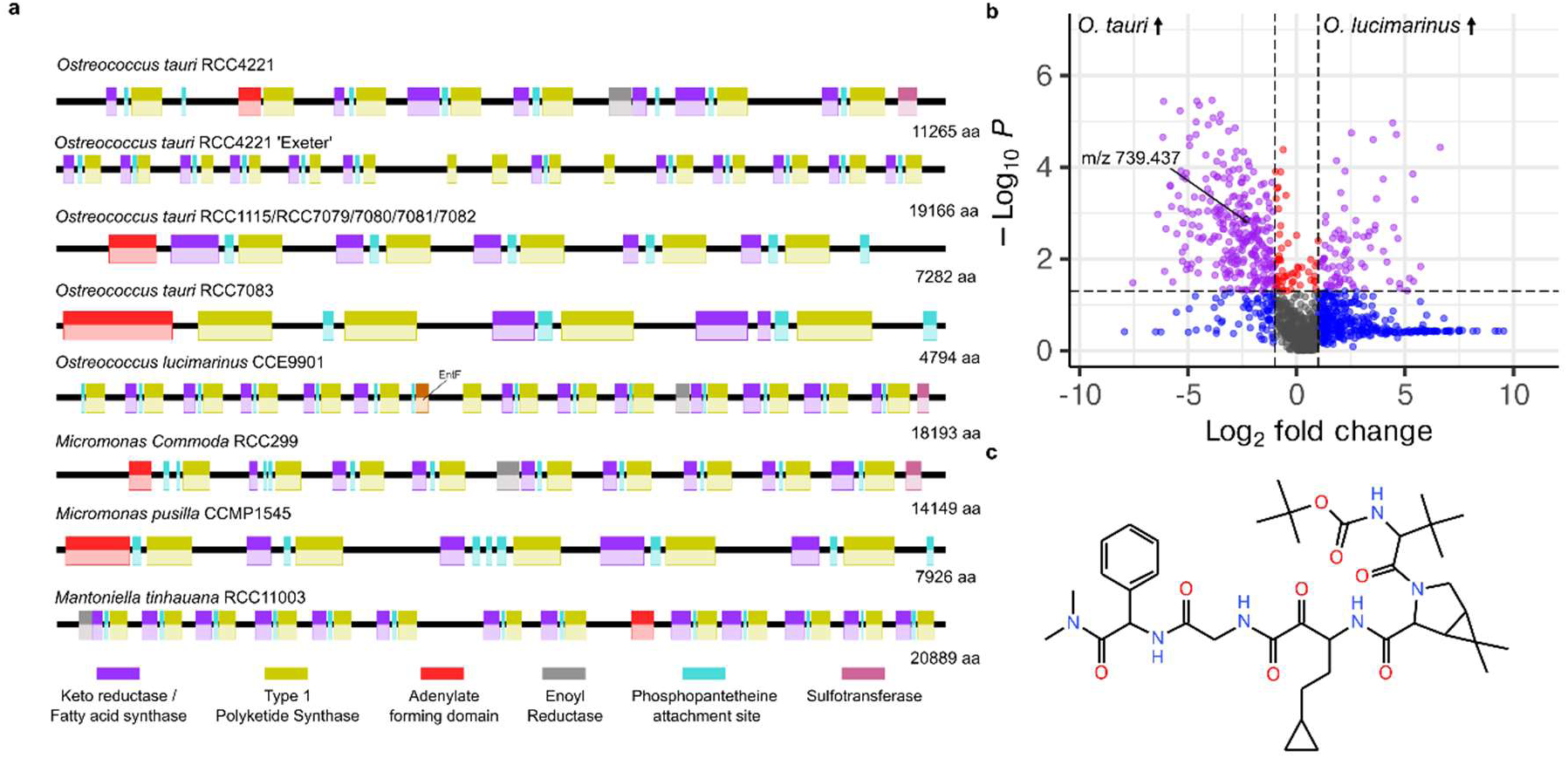
Modular Giant Type I Polyketide synthases show structural diversity across Mamiellales species. (**a**), Schematic representation of the largest orthologous polyketide synthases identified across Mamiellales species, illustrating large size variation in protein product, from 4,794 amino acid residues in *O. tauri* RCC7083 to 20,889 residues in *M. tinhauana* driven by differences in modular domain repetition. Coloured boxes represent different functional domains within the modular PKS architecture. (**b**), Volcano plot showing differential metabolite abundances between *O. lucimarinus* (CCE9901) and *O. tauri* (RCC4221) across 1,318 unidentified positive ion features, revealing 273 features enriched in *O. tauri* versus 81 enriched in *O. lucimarinus*. Fold-change thresholds and statistical significance (p < 0.05) are indicated by dashed lines. A putative polyketide showed 5.29-fold higher abundance in *O. tauri* compared to *O. lucimarinus* (p = 0.0016, ANOVA; RT 14.38 min, m/z 739.437, ID#3563). Gray dots represent non-significantly different metabolites (see Dataset S16–18 for additional structural and metabolomic analyses). (**c**), Structure of the putative polyketide-like compound inferred from LC–MS/MS fragmentation patterns.

To explore whether these structural variations correspond to metabolic divergence, we performed untargeted metabolomics on *O. tauri* (RCC4221) and *O. lucimarinus* (CCE9901), two species with contrasting PKS organisations in BOC1 (SI Appendix, Fig. S13; Dataset S15). Across both ionisation modes, we detected 2,213 unknown mass features, with 273 and 50 showing significantly higher abundance in *O. tauri* and 81 and 41 in *O. lucimarinus* in positive and negative ion modes, respectively (ANOVA, p<0.05, |Log2FC|>2; Fig. 6b; Dataset S16 and S17). These comparisons reveal broad chemical divergence between the species. Among the discriminant features, we identified one compound (RT 14.38 min, m/z 739.437; Fig. 6c; SI Appendix, Fig. S14; Dataset S18) whose fragmentation pattern is consistent with polyketide-like structures, though structural confirmation and direct demonstration of PKS involvement remain to be established.

### Similar outlier architecture in phylogenetically distant marine picoeukaryotes

To determine whether spatially insulated outlier regions occur in other minimal marine eukaryotes, we analysed genomic data from diverse lineages, including those sequenced by the Darwin Tree of Life project (29). In the stramenopile *Pelagomonas calceolata* (30), we identified discrete ∼500 kb outlier regions on each chromosome characterised by ∼5% lower GC content and significantly elevated TE density (Mann-Whitney U test p<0.0001; SI Appendix, Fig. S15). Analysis of available Hi-C data confirmed that these regions are spatially compartmentalised from the standard genome (30), exhibiting insulation architecture similar to that observed in Mamiellales BOC1 and SOC1. Genes within *P. calceolata* outlier regions show ∼3-fold elevated expression compared to standard regions. Remarkably, one *P. calceolata* outlier region contains a giant PKS gene positioned adjacent to a MULE element (SI Appendix, Fig. S16). We also identified similar outlier features in the diatom *Epithemia pelagica* (Bacillariophyceae) (31), where specific chromosomes (e.g., OX337238.1) harbour dense, localised clusters of TEs, predominantly LTR retrotransposons of the Gypsy superfamily (97 copies in the outlier region; Dataset S19).

We next examined other Chlorophyta classes. In the Pseudoscourfieldiophyceae, we analysed the genomes of *Pycnococcus provasolii* and *Pseudoscourfieldia marina* (32). Although historically classified as distinct genera, recent organellar phylogenomics has unified them as morphotypes of the single species *P. marina* (32). Consistent with this reclassification, we observed 98% average nucleotide identity between the two assemblies and high synteny across their standard chromosomes. However, we identified distinct, low-GC outlier regions in both genomes (Mann-Whitney U test, p<0.0001; Dataset S19). While the standard chromosomes are highly conserved, the outlier regions exhibit marked structural divergence and TE load differences between the two morphotypes (SI Appendix, Fig. S17), paralleling the intraspecific divergence observed in *Ostreococcus*. Finally, in the trebouxiophyte *Marvania coccoides* (33), we identified a single chromosome (OY978099.1) containing a discrete region with massive TE accumulation (362 elements, including 106 LTR retrotransposons, compared to a genome average of <50 per comparable window; Mann-Whitney U test p<0.0001; Dataset S19).

The recurrence of spatially defined, TE-rich, low-GC compartments across the chlorophytes (Mamiellophyceae, Pseudoscourfieldiophyceae, and Trebouxiophyceae), and stramenopiles (Pelagophyceae and Bacillariophycea), indicates that this chromosomal organisation has arisen independently in phylogenetically distant lineages of marine eukaryotes.

## Discussion

### Outlier regions form spatially insulated compartments with distinctive chromatin properties

Mamiellales, among the smallest free-living eukaryotes, harbour unusually large and dynamic chromosomal regions (BOC1 and SOC1) that represent distinct genomic compartments. Our multi-omics analysis reveals that these outlier regions share a conserved suite of structural features across species: reduced DNA methylation, elevated gene expression, and spatial insulation from the rest of the genome. The Hi-C data presented here demonstrate that BOC1 and SOC1 form TAD-like compartments, with sharp boundaries that align precisely with epigenetic transitions and changes in TE density.

The transcriptional hyperactivity of genes within outlier regions mirrors patterns observed in low-GC regions of land plants such as *Oryza sativa* (34), but contrasts with the freshwater alga *Chlamydomonas reinhardtii*, where low GC content correlates with reduced expression (35). This contrast suggests that marine picoeukaryotes and their freshwater relatives have evolved distinct regulatory architectures governing gene expression in low-GC regions. The hypomethylated state of outlier regions, combined with their spatial insulation, is associated with a chromatin environment that is transcriptionally permissive and physically separated from the compact, gene-dense standard genome.

### Nucleosome organisation is structured and may influence TE distribution

Contrary to earlier cryo-electron tomography studies suggesting disordered chromatin in *O. tauri* (36, 37), our re-analysis of MNase-seq data (27) reveals that chromatin organisation in these minimal eukaryotes is structured and species-specific. The outlier regions in *O. lucimarinus* and *M. pusilla* display well-phased, closely spaced nucleosomes, which is a configuration associated with chromatin flexibility (38, 39). In *M. pusilla*, the bimodal distribution of TEs around nucleosome centres suggests that chromatin structure may influence TE insertion patterns, with elements preferentially integrating into accessible linker DNA. This parallels findings in *Arabidopsis*, where the histone variant H2A.Z influences TE insertion site preference (40). Given the presence of plant-like H2A.Z histone variants, which are known to influence chromatin structure and TE insertion, in Mamiellales, similar mechanisms may contribute to the concentration of TEs within outlier regions.

### Three-dimensional organisation, not TE content, defines outlier regions

A key finding of our study is that several features of outlier regions are conserved across Mamiellales, regardless of TE content. This is most clearly illustrated by *O. lucimarinus* and *B. prasinos*, which lacks substantial TE enrichment yet retains the characteristic low-GC profile and elevated gene expression observed in TE-rich species. While Hi-C data is unavailable for both TE-depleted species, the persistence of these hallmark features suggests that the underlying genomic context, likely involving conserved chromatin regulation, is maintained independently of TE accumulation. This argues against models in which TEs are the primary drivers of outlier region properties, instead suggesting that TEs accumulate as a consequence of the unique genomic environment of these regions.

### Giant PKS genes exhibit extensive structural variation

The giant type I PKS genes found in BOC1 represent one of the most striking examples of structural variation within these regions. PKS enzymes synthesise diverse secondary metabolites including toxins, antibiotics, and signalling molecules that can be important for ecological interactions (41, 42). We found that MULEs frequently flank or insert within PKS loci. Since MULEs are known to mediate gene duplication and shuffling (43), their proximity to PKS genes is notable, though whether they actively contribute to PKS diversification remains to be determined. Our long-read sequencing revealed substantial structural variation within the PKS locus of *O. tauri* RCC4221 compared to the reference genome, and untargeted metabolomics detected species-specific chemical differences between *O. tauri* and *O. lucimarinus*. Among these, one feature showed properties compatible with a polyketide-like compound, though its structure and biosynthetic origin remain unconfirmed. Establishing a direct link between PKS gene variation and metabolic output will require targeted biochemical characterisation.

### Structural parallels with sex-determining regions

The structural features of BOC1, suppressed recombination and TE accumulation, resemble the sex-determining regions of other eukaryotes (44, 45). This organisation parallels the UV sex chromosome systems of brown algae such as *Ectocarpus*, where non-recombining regions accumulate TEs and lineage-specific genes (46, 47). Recent Hi-C analysis of *Ectocarpus* confirms that its sex-determining regions are spatially insulated from flanking pseudoautosomal regions (48), directly paralleling the TAD-like compartmentalisation we observe in Mamiellales. However, an important distinction exists: while *Ectocarpus* pseudoautosomal regions often show lower expression (47), the *Ostreococcus* outlier regions are transcriptionally hyperactive. Whether the outlier regions of Mamiellales evolved from ancestral mating-type loci, or whether the similarities reflect convergent structural solutions to distinct biological problems, remains an open question.

### Convergent genome compartmentalisation across marine picoeukaryotes

The identification of similar outlier architecture in phylogenetically distant lineages is perhaps the most unexpected finding of our comparative analysis. In the stramenopile *P. calceolata*, we found discrete low-GC regions with elevated TE density, spatial insulation (30), increased gene expression, and, remarkably, giant PKS genes adjacent to MULE elements. Similar features occur in the diatom *E. pelagica* (31) and in chlorophyte lineages outside the Mamiellophyceae. The recurrence of this chromosomal organisation across Mamiellophyceae, Pseudoscourfieldiophyceae, Trebouxiophyceae, Pelagophyceae, and Bacillariophyceae suggests either deep ancestry predating the divergence of these groups, or, more likely given the phylogenetic distances involved, independent origins of similar genome organisation. What selective pressures or mechanistic constraints might favour this arrangement remains unclear. One possibility is that spatial insulation provides a means to tolerate TE activity without disrupting essential genes; another is that the chromatin environment of these regions is permissive for the expression of large, complex genes like PKS loci that might be poorly expressed elsewhere in compact genomes. These hypotheses are not mutually exclusive and remain to be tested.

### Tolerance without silencing: an unconventional epigenetic architecture

Our findings contribute to an emerging picture of how genome architecture varies across diverse eukaryotes. Similar patterns of compartmentalised genomic divergence have recently been reported in bivalve mollusks, where low-GC, TE-rich “Haplotype Divergent Sequences” show elevated structural variation (49). In contrast to many eukaryotes, that silence repetitive regions via hypermethylation (50, 51), the outlier regions of minimal marine eukaryotes maintain a hypomethylated, transcriptionally active state. Whether this reflects distinct regulatory mechanisms, weaker silencing machinery, or active maintenance of an “open” chromatin state remains to be determined. Key questions for future work include the molecular mechanisms that establish and maintain the sharp boundaries between outlier and standard regions, whether outlier chromosome polymorphisms contribute to phenotypic traits such as viral susceptibility (17, 18), and the evolutionary history of these regions across eukaryotic lineages. Our analysis establishes that outlier chromosomes are spatially insulated compartments defined primarily by their three-dimensional organisation, rather than by specific sequence composition. The independent occurrence of similar chromosomal architecture in distantly related marine eukaryotes points to shared fundamental principles, whether adaptive or mechanistic, that govern genome organisation in compact eukaryotic genomes.

## Materials and Methods

### Algal strains and culturing

A total of eight Mamiellales species were included in the study, representing the genera *Ostreococcus*, *Bathycoccus*, *Micromonas* and *Mantoniella*. We used *Ostreococcus tauri* RCC4221 in this study to generate data for Hi-C, RNA-Seq and methylation profiling. We sourced other molecular datasets from publicly available databases. The species, strain IDs and datasets used in this study are provided in SI Appendix, Table S1.

*O. tauri* strain RCC4221 was maintained in L1 seawater medium (52) in non-treated polycarbonate flasks and maintained in a 14 h: 10 h light: dark cycle at 35 µmol m^−2^ s^−1^ irradiance and a temperature of 20 °C. To achieve a near-axenic culture of *O. tauri*, we isolated cells by flow cytometric cell sorting (BD Influx, Beckton Dickinson) and maintained algal cells in exponential growth by weekly subculturing. We also used a monthly antibiotic treatment (5 µg Ampicillin, 1 µg Gentamicin, 2 µg Kanamycin, 10 µg Neomycin). Algal growth was regularly monitored by flow cytometry (Cytoflex, Beckman Coulter) based on side-angle light scatter and chlorophyll red fluorescence. Bacterial cells were monitored and quantified via flow cytometry after SYBR Green I staining.

### Transcriptomics

#### RNA extraction and sequencing

*O. tauri* RCC4221 cells were collected by centrifuging a 20 mL sample (10 min at 4000 × g at 4 °C), snap-frozen in liquid nitrogen, and stored at −80 °C. RNA was extracted using a RNeasy Plant kit (QIAGEN) with RLT buffer incubated at 56 °C for 3 min. The residual DNA was digested using the Turbo DNA-free kit (Thermo Fisher Scientific). RNA was quantified using a Qubit Fluorometer (Thermo Fisher Scientific), and RNA quality was assessed using an Agilent 2100 Bioanalyzer System. RNA extracts with an RNA integrity number (RIN) greater than eight were subjected to poly(A) selection prior to library preparation with the Illumina TruSeq Direction Library Prep Kit and sequenced on Illumina NovaSeq 6000 (150 bp paired end reads).

#### RNA-Seq data processing

Adapters and low-quality reads from RNA-Seq libraries were trimmed using Fastp v0.23.2 (53) (minimum length required 50). Quality control reports were compiled using FastQC v0.11.9 (54), MultiQC v1.13 (55), and fastp. Ribosomal RNA reads were removed using SortMERNA v4.3.6 (56) matching reads with rRNA sequences from the SILVA databases (57). The remaining reads were aligned and quantified to the *O. tauri* reference genome sequence (GCF_000214015.3) using Star v2.7.10 (58) with default parameters. Counts were divided by gene length to standardise counts and generate transcript levels. For publicly available RNA-Seq libraries and counts, transcript levels were measured in *M. pusilla* CCMP1545 using table ‘Sup_Table_S2_CCMP1545’ column ‘P-Replete T1.5 hours’ for counts (59). For *M. commoda* RCC299, existing triplicate axenic counts were used (60). For *B. prasinos* RCC1105, *O. lucimarinus* CCE9901, *M. commoda* RCC299, the reads from SRR1300453 (61), SRR847308 (27) and SRR17220685 (60) were used with the same workflow as above to additionally measure splice events with Star.

### Oxford Nanopore Technology sequencing

#### DNA extraction and library preparation

Cells were collected by centrifuging a 30 mL sample (10 min at 4000 × g at 4 °C), snap-frozen in liquid nitrogen, and stored at -80 °C. DNA was extracted using a DNeasy Plant kit (QIAGEN). DNA was quantified using a Qubit Fluorometer (Thermo Fisher Scientific). Library preparation was conducted using the Ligation Sequencing Kits (SQK-LSK109; SQK-LSK114; ONT, Oxford, UK). Genomic DNA was pooled for sequencing on an FLO-FLG001 Flongle flow cell and FLO-MIN114 Minion flow cell (R9.4.1; R10.4.1; ONT, Oxford, UK).

#### Methylation profiling

Super-accuracy base calling was performed on R10.4.1 sequence data using Dorado v0.6.0 (ONT, Oxford, UK), Methylation profiling included N6-methyladenine (6mA), N5-methylcytosine (5mC/5mCG), and N5-hydroxymethylcytosine (5hmC/5hmCG) which were transformed into BED files using modkit v0.2.8 (ONT, Oxford, UK). To normalise for variations in local GC content, % methylation levels represent the average proportion of the methylated base relative to the total number of that base within the sliding window.

#### Genome assembly

Exeter RCC4221 was de novo assembled from R9.4.1 ONT sequence data using Flye v2.9.2 (62) (--iterations 3 –scaffold) followed by medaka_consensus v1.8.0 (ONT, Oxford, UK). Matching scaffolds to reference were pulled out and aligned using blastn v2.1.0 (63).

### High-throughput chromosome conformation capture

#### Hi-C sequencing

For consistency, samples for all Hi-C experiments were collected at 2 hours into the light phase. To ensure sufficient DNA yield for sequencing from small *O. tauri* genomes, 1–3 × 10^8^ total cells were used in all experiments. As there was no published protocol for picoeukaryotes we tested both *in situ* (64) and *in vivo* Hi-C approaches and a series of optimisation steps, which are detailed in the SI Appendix. In summary, the final protocol was as follows: we performed crosslinking by adding 2% v/v final concentration formaldehyde (16% TEM grade, Sigma) directly within the culture medium containing the cells and incubating at room temperature for 20 mins. Cross-linking was stopped by adding glycine (0.2 M final concentration) and incubating for 5 mins at room temperature on a roller. Finally, cells were washed with ice-cold PBS and stored at −80 °C. The final Hi-C library was generated using the Proximo Microbe Hi-C kit v2 (PhaseGenomics, USA) following the manufacturer’s instructions and sequenced using the Illumina NovaSeq 6000 platform (150 bp paired end reads) yielding a total of 94143446 reads (run ID V0017_combined in Dataset S10).

#### Hi-C data processing

Adapters and low-quality reads were trimmed from Hi-C libraries using Fastp v0.23.2. Quality control reports were compiled using FastQC v0.11.9 (54), MultiQC v1.13 (55), and fastp. The Hi-C processing pipeline, with the restriction sites of “^GATC,^AATT” and ligation sites of “GATCGATC,GATCAATT,AATTGATC,AATTAATT”, was based on NextFlow v22.04.5 (available as a Docker profile: nf-core/hic v1.3.0) (65–67). Hi-C reads were mapped onto a soft and hard-masked (SI Appendix, Fig. S18) *O. tauri* reference genome sequence (GenBank identifier: GCF_000214015.3) using Bowtie v2.3.5.1 (68). Duplication removal and contact pair validation were conducted with HiC-Pro v2.11.1 (65). Raw and normalised contact maps were generated using cooltools v0.3.0 (69) at bin resolutions of 5, 10, and 20 kb. Additional normalisation was carried out using ‘iced’ with HiC-Pro, a Python implementation of the ICE normalisation algorithm from Cooltools (70). Hi-C quality reports were generated using HiCUP (71). Domain compartment calling was completed using Cooltools. The R package HiCRep (72) was used to evaluate the correlation between Hi-C experimental replicates (SI Appendix, Fig. S19). Juicer v1.6 (73) was used to produce additional contact maps for use with downstream loop detection. Chromatin loops were identified from the Nextflow and Juicer contact maps using Mustache v1.0.1 (74) and HiCCUPS v3.0 (73, 74), respectively, at bin resolutions of 5, 10, and 20 kb. The derivative/slope of contact probability, cis, and trans interactions was calculated using cooltools. *M. tinhauana* RCC11003 Hi-C data was similarly analysed (25).

### Additional sequence analyses

#### MNase-seq analysis

MNase-seq data for *M. pusilla* CCMP1545 and *O. lucimarinus* CCE9901 (SRR847304 and SRR847307) (27) were processed to analyse nucleosome positioning. Paired-end reads underwent quality trimming and adapter removal using fastp v0.23.4. The cleaned reads were aligned to the reference genome with Minimap2 v2.24 (75) using default parameters for paired-end data. Aligned files were converted to BAM format, sorted, and indexed with Samtools v1.13 (76). Finally, genome-wide coverage was calculated and normalised to RPKM using bamCoverage (deeptools v3.5.1 (77), --MNase --center-reads) to calculate nucleosome centres, and the results were exported as bigwig files for visualisation and downstream analysis in Python for peak calling. Autocorrelation functions of nucleosome center coverage were estimated using python. Nucleosome fuzziness and stiffness were calculated using Nucleosome Dynamics v1.0 (78).

#### Bisulfite-seq analysis

Bisulfite sequencing data for *M. pusilla* CCMP1545 and *O. lucimarinus* CCE9901 (SRR847303 and SRR847306 (27)) were processed to determine methylation levels at CpG sites. Single-end reads were trimmed for quality and adapters using Fastp v0.23.4. The trimmed reads were aligned to the reference genome using Bismark v0.24.2 (79). Methylation calling was restricted to CpG contexts, and methylation levels were summarised in bedgraph and bigwig format for further analysis.

#### Homology and GO analyses

Protein synteny was highlighted using Oxford dot plots v0.3.3 (80), blastp v2.10 (63) and Diamond v2.1.8 (81) across outlier chromosomes in the Mamiellophyceae. A ribbon plot was generated to show the synteny between species using Python. Updated protein annotations were determined using AlphaFold3 (82) and FoldSeek (83). EDTA v2.2.2 (84), default settings, were used to flag transposable elements across whole genomes and PKS genes, respectively. Enrichment of TEs was calculated using 5 kb bins and Kolmogorov–Smirnov test. A data frame of all the proteins within *O. tauri* with GO enrichment spanning biological processes, molecular functions, cellular components, and locations on chromosomes was obtained from NCBI and UniProt (85). Fisher’s exact test (one-tailed) was performed using a custom script in Python on subsets of genes to compare enriched terms over target regions and the rest of the genome.

### Untargeted metabolomics

Metabolite profiling was carried out using LC-quadrupole time of flight mass spectrometry with separate runs for positive and negative ion modes. *O. tauri* RCC4221 and *O. lucimarinus* CCE9901 cells were collected by centrifuging a 30 mL sample (10 min at 4000 × g at 4 °C), snap-frozen in liquid nitrogen, and stored at −80 °C. Pellets were extracted in 0.6 mL methanol, treated for 20 min in a sonicating water bath with ice and then centrifuged at 20,000 × g for 10 min at 4 °C. The supernatants from three replicate cultures of each species and extraction blanks were analysed using an Agilent Zorbax Eclipse plus, narrow bore RR, 2.1 × 50 mm, 3.5 μm particle size reverse phase C18 analytical column (Agilent Technologies, Palo Alto, USA). 5 μL samples were injected *via* an Agilent 1200 series Rapid Resolution HPLC system coupled to a quadrupole time-of-flight (QToF) 6520 mass spectrometer (Agilent Technologies, Palo Alto, USA). Mobile phases were A (10% methanol, 0.1% formic acid), mobile phase B (100% methanol with 0.1% formic acid). The following gradient was used: 0–5 min-30% B; 10–24 min-0–98% B; 25-30 min-100% B. The flow rate was 0.3 mL min^−1^ and the column temperature was 40 °C. The source conditions for electrospray ionisation were as follows: gas temperature was 325 °C with a drying gas flow rate of 9 L min^−1^ and a nebuliser pressure of 35 psig. The capillary voltage was 3.5 kV in both positive and negative ion mode. The fragmentor voltage was 175 V and skimmer 65 V. Scanning was performed using the auto MS/MS function at 5 scans s^−1^ for precursor ion surveying and 4 scans s^−1^ for MS/MS with a sloped collision energy of 3.5 V/100 Da with an offset of 10 V. MS and MS/MS m/z range was 40–1700. Reference compound m/z values were 121.05087 and 922.00979. Raw data files (*.d) were converted to mzML format using MSConvert (ProteoWizard) (86). Feature detection and alignment was carried out with MS-Dial v5 (87). Features present in the blank (>20% of sample intensity) were discounted. Compounds were identified with the MS-Dial lipidomics workflow and using publicly available CompMS MS-Dial MS/MS libraries. Statistical one-factor Anova test analysis (p<0.05, |LFC|>2) was then performed on ‘Unknown’ features with Metaboanalyst v6.0 (88) to highlight putative PKS products for further annotation within Sirius (89).

## Data availability

Sequencing data have been deposited at NCBI under BioProject accession PRJNA1250464. Additional data including Hi-C contact matrices and metabolomic profiles are available at Zenodo: 10.5281/zenodo.14284467.

## Acknowledgements

Illumina sequencing was conducted on equipment funded by the Wellcome Trust (Multi-User Equipment Grant award number 218247/Z/19/Z). We are grateful to Dr Karen Moore (Exeter Sequencing Facility, University of Exeter) for their technical advice on Hi-C sequencing. We thank Dr Aurélie Chambouvet and Dr Evelyne Derelle (CNRS, France) for providing algal strains. This research was supported by the Royal Society through a University Research Fellowship (URF\R1\180537) and Research Fellows Enhancement Awards (RGF\EA\181012) awarded to AM.

