## Supplementary Information for "Spatial Insulation Confines Dynamic Chromosomes in Minimal Marine Eukaryotes"

**Competing Interest Statement:** No competing interests

**Keywords:** Chromosome conformation capture, Genomic compartmentalisation, picoeukaryotes, transposable elements, polyketide synthase.

**This PDF file includes:**

Supporting Information Text (Extended Methods)
Figures S1 to S18
Tables S1 to S4
Legends for Datasets S1 to S19
SI References

**Other supporting materials for this manuscript include the following:**

Datasets S1 to S19

### Supporting Information Text

### Extended Methods

#### High-throughput chromosome conformation capture (Hi-C) protocol optimization

To generate high-quality Hi-C maps for O. tauri, we tested both an *in situ* Hi-C approach originally intended for metazoans (4DN Nucleome in-situ Hi-C protocol), as well as *in vivo* Hi-C using the Proximo Microbe kit (Phasegenomics, USA). Several optimization steps were necessary due to the small size of the cells, compact genomes, and the presence of chlorophyll and cellulose,

***Cell collection and DNA cross-linking.***

Most of the optimisation was necessary for DNA cross-linking and reverse cross-linking of DNA, considering that small cells are easily damaged by centrifugation and that high polysaccharide and chlorophyll content interferes with downstream processing. Specifically, we found that the high centrifugation speed required to concentrate small *O. tauri* cells increased the number of interchromosomal contacts and masked intrachromosomal features, possibly due to the deformation or destruction of some cells. Therefore, we performed crosslinking by adding formaldehyde directly on cells within the culture medium. Based on the literature, we tested 1%, 2% and 3% v/v final concentration of formaldehyde (16% TEM grade, Sigma) and 10 versus 20 minutes incubation time at room temperature on a roller. The best combination for recovering a higher number of intrachromosomal contacts whilst not overfixing the small, compact cells was 2% formaldehyde for 20 mins (Dataset S8). Cross-linking was stopped by adding glycine (0.2 M final concentration) and incubating for 5 mins at room temperature on a roller. Finally, cells were washed with ice-cold PBS and then stored at -80 °C until further processing.

***In-situ Hi-C.*** Initial Hi-C runs were attempted using the 4DN Nucleome in-situ Hi-C protocol (https://www.4dnucleome.org/protocols.html). As this protocol was originally developed for metazoan cells, several modifications were necessary because of the small nuclei and genome, presence of chlorophyll, and large amounts of polysaccharides in *O. tauri*. Cell lysis and nuclei isolation were performed by 4 x 10 min incubations in ice-cold lysis buffer (10 mM Tris-HCl pH 8.0, 10 mM NaCl, 0.2% v/v Igepal and 1x complete, EDTA-free, protease inhibitor cocktail). Changing the lysis buffer every 10 mins was a crucial step to remove cell debris and chlorophyll and efficiently isolate pure nuclei. Nuclei were permeabilized by incubating at 62 °C for 10 min in 0.5% SDS and 37 °C for 15 min in 0.5% Triton X-100. Chromatin was fragmented using the restriction enzyme 100 U Dpn II (restriction site GATC) at 37 °C for 1 h 30 min and ends were filled in with nucleotides including biotinylated dATP. Proximity ligation was performed overnight at 16 °C. Chromatin was treated with RNAse A and cross-links were reversed by overnight (maximum 16 hours) incubation with Proteinase K (3.3 mg/mL final concentration) at 68 °C; this also released DNA from the nuclei. DNA was isolated using ice-cold isopropanol for 1 hour at -20 °C followed by 30 mins centrifugation at maximum speed. DNA pellets were washed twice with 75% Ethanol and resuspended in 10 mM Tris buffer. The Hi-C libraries were visually verified by agarose gel electrophoresis. DNA was sheared using a Covaris instrument to approximately 300 bp fragments, and biotinylated fragments were selected using streptavidin magnetic beads (Invitrogen, Waltham, United States). The library was end repaired, A-tailed and adapter-ligated using the NEB Next Ultra II FS kit and amplified with NEB Next dual index barcodes for sequencing in the Illumina MiSeq platform.

***Proximo Microbe Hi-C kit.*** Due to excessive interchromosomal contacts retrieved with the in situ Hi-C protocol even after many trials and modifications of the protocol, we tested the commercial Proximo Hi-C Microbe kit by PhaseGenomics (USA), which is especially formulated for microbial organisms. The manufacturer's protocol was followed post-DNA-crosslinking, which included complete cell lysis followed by immobilisation of chromatin onto magnetic beads. For fragmentation, the kit contains two enzymes, Sau3AI which is equivalent to DpnII but with different methylation sensitivity (restriction site GATC), and MluCI (restriction site AATT). Biotinylation, proximity ligation, reverse cross-linking, enrichment of biotinylated fragments, and library preparation for Illumina sequencing were performed on the immobilised DNA using the kit's custom reagents.

***Hi-C sequencing.*** To compare the two protocols, duplicate samples were collected and subjected to Hi-C. Libraries were quality assessed for the correct size by electrophoresis and amplification efficiency by qPCR. Libraries were sequenced on a MiSeq Illumina platform. We obtained 13,417,158 and 94,143,446 150bp paired end reads per sample for in situ and Proximo Hi-C, respectively. The difference in the number of reads was because of low library concentrations obtained with the Hi-C protocols. A comparison of the two approaches showed that the commercial kit was superior in recovering intrachromosomal interactions, while in situ Hi-C sequencing data were dominated by potentially random interchromosomal interactions. The ratio of intra- versus inter- chromosomal interactions was 3.43 and 0.288, respectively, for the two protocols (Dataset S8).

### Figures

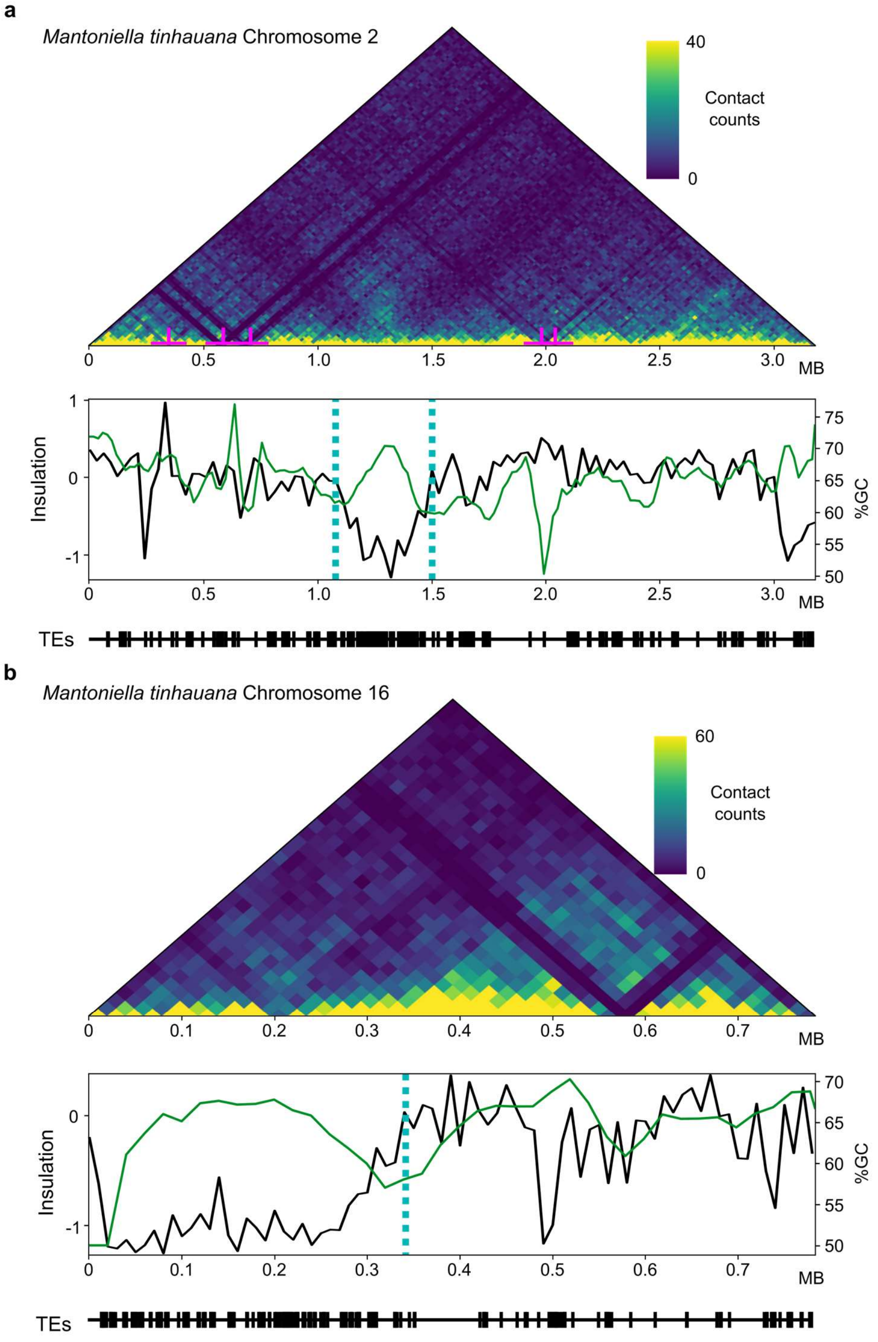


**Figure S1. Chromosomal organisation of BOC and SOC within Mantoniella tinhauana.** (**a—b**), Hi-C contact maps binned at 10 kb with normalised counts for chromosome 2 and chromosome 16. Below tracks include %GC-content in black and insulation score in green across the chromosome. Alongside TE occurence across the chromosome.

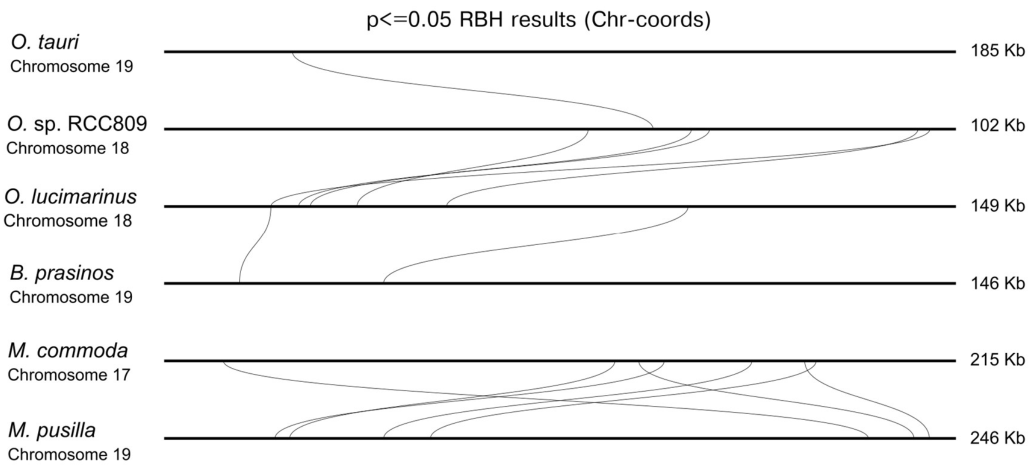


**Figure S2. Conservation, synteny and functional enrichment of Mamiellales SOC1 regions.** Homology plots highlighting SOC1 regions across outlier chromosomes in seven Mamiellales species. Each horizontal line represents a chromosome, with the species name and chromosome number indicated on the left. The length of each chromosome is shown in megabases (Mb) on the right. Green areas represent the SOC1. Lines connecting the chromosomes represent homology relationships based on reciprocal best BLAST hits.

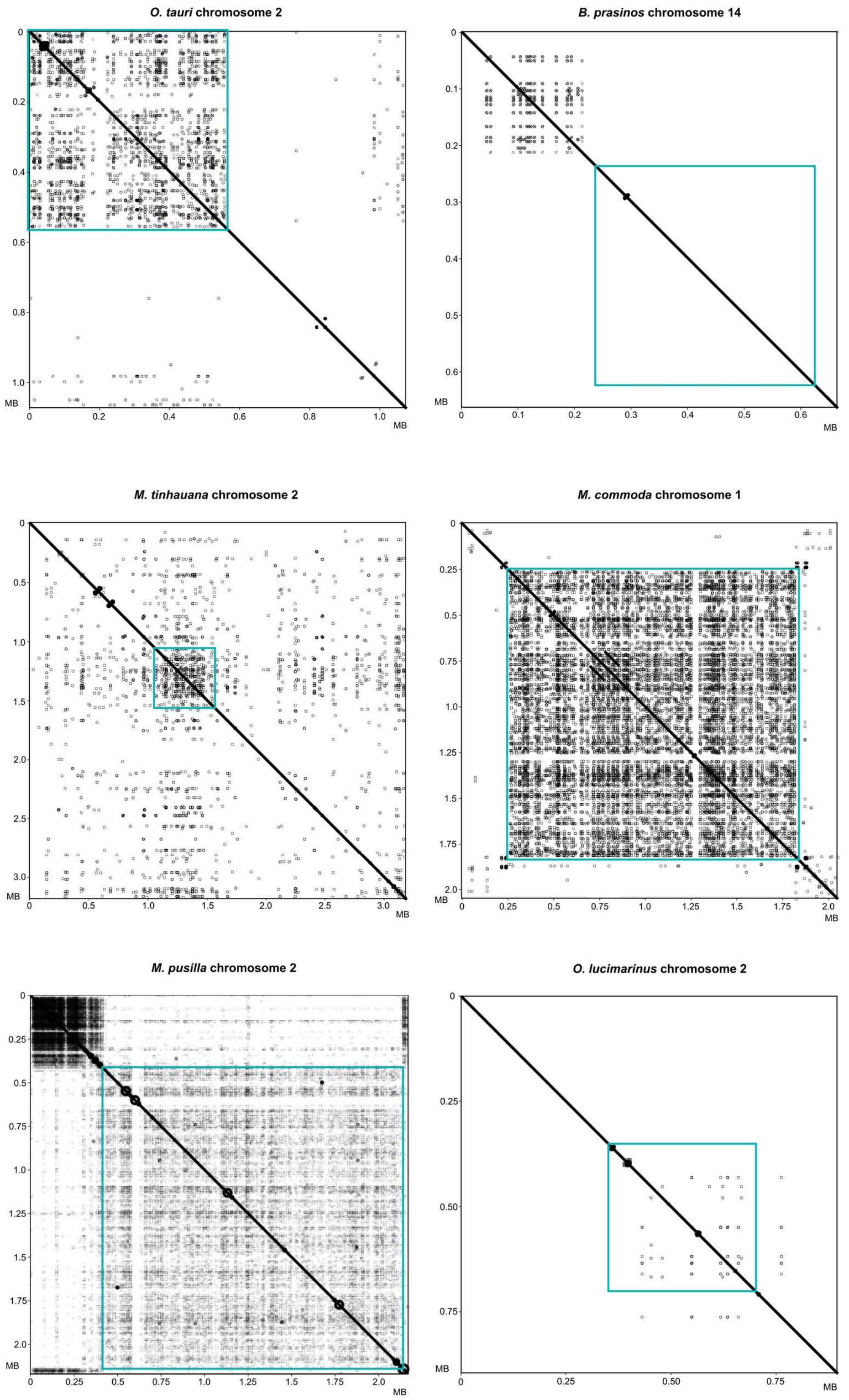


**Figure S3. High nucleotide similarity of BOC1s.** Dot plots of reciprocal blastn results of BOCs across Mamiellales. Blue squares indicate BOC1 region.

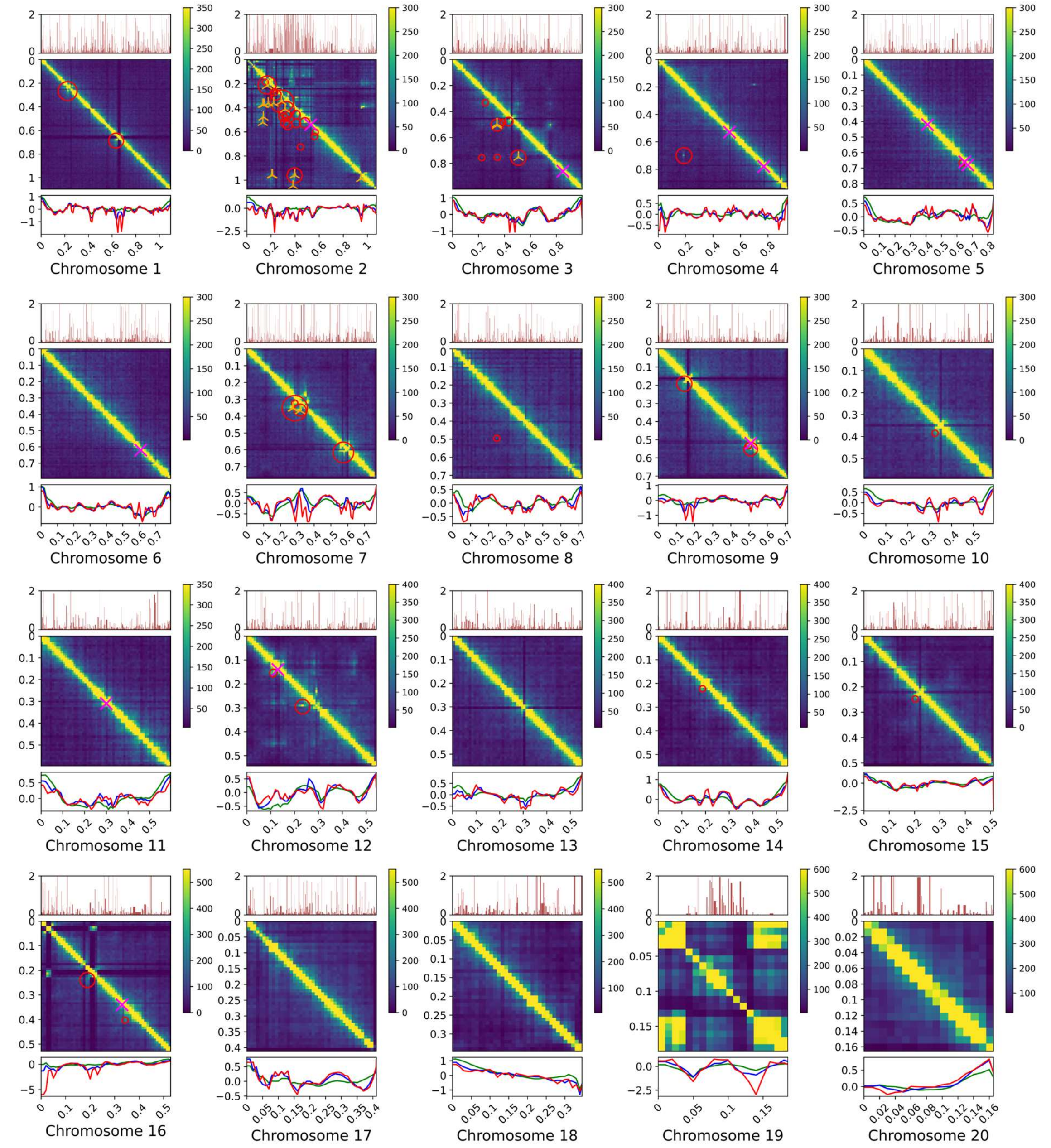


**Figure S4. Hi-C contact maps, transcription, insulation scores for all chromosomes in *O. tauri*.** Hi-C contact maps, 10 kb resolution for each nuclear *O. tauri* chromosome. Chromatin loops identified using Hiccups and Mustache are coloured using orange crosses and red circles, respectively. Topologically associating domains are represented using fuchsia crosses. The inner and outer upper tracks represent the rolling average of a scaled window size of the proportion of 5mCG (orange), 5mC (green), 6mA (purple), 5hmCG (red), 5hmC (blue) modifications, %GC-content, and transcript levels across the chromosome, respectively, whilst the lower track is the insulation score, the sum of Hi-C contacts in a small sliding window.

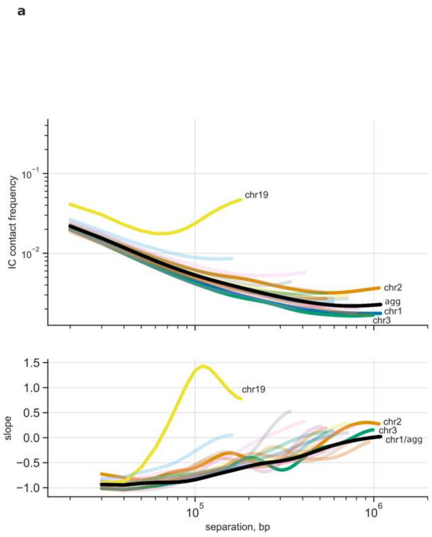


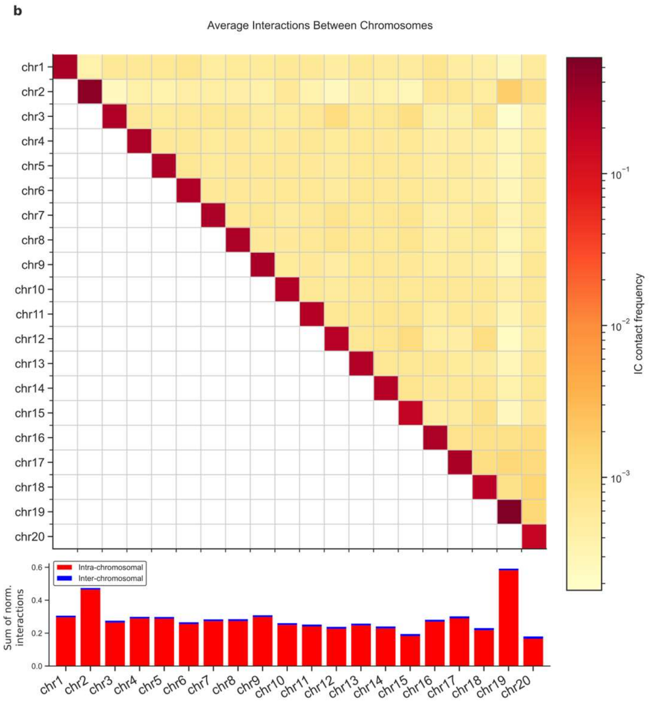


**Figure S5. Inter- and Intra-chromosomal contacts by chromosome.** (**a**), Dependence of interaction probability on genomic distance smoothed across each chromosome alongside an aggregation of all chromosomes (shown in previous figure) alongside its derivative with respect to separation (bp). (**b**), Heatmap of normalised inter-chromosomal contact frequency across each chromosome with a bar chart showing the sum of inter- and intra-chromosomal contacts.

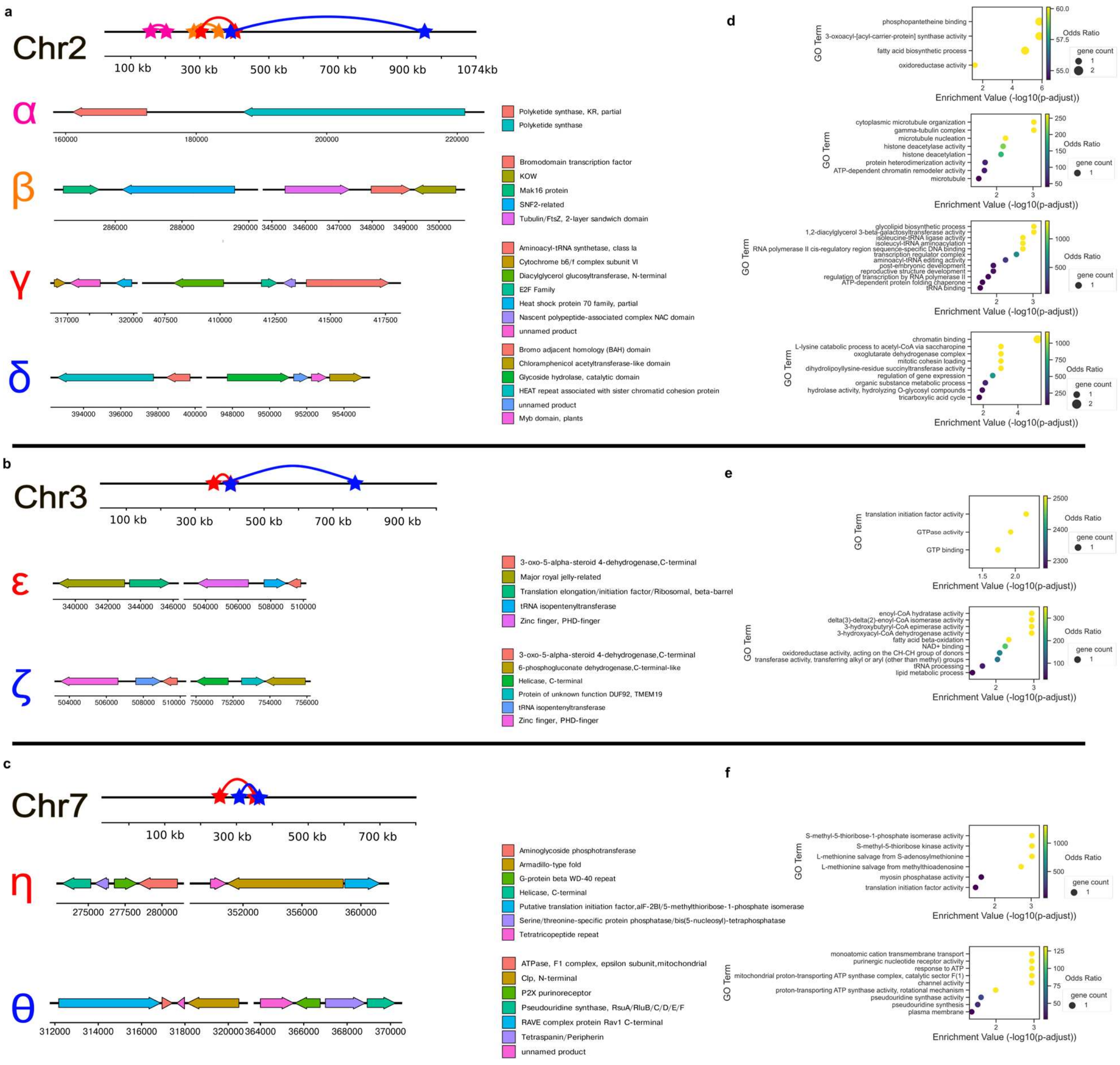


**Figure S6. Occurrence of genes across chromatin loops**. (**a**), Genes on chromosome 2 chromatin loops. (**b**), Genes on chromosome 3 chromatin loops. (**c**), Genes on chromosome 7 chromatin loops. (**d**), GO Enrichment for genes in **a**. (**e**), GO Enrichment for genes in **b**. (**f**), GO Enrichment for genes in **c**.

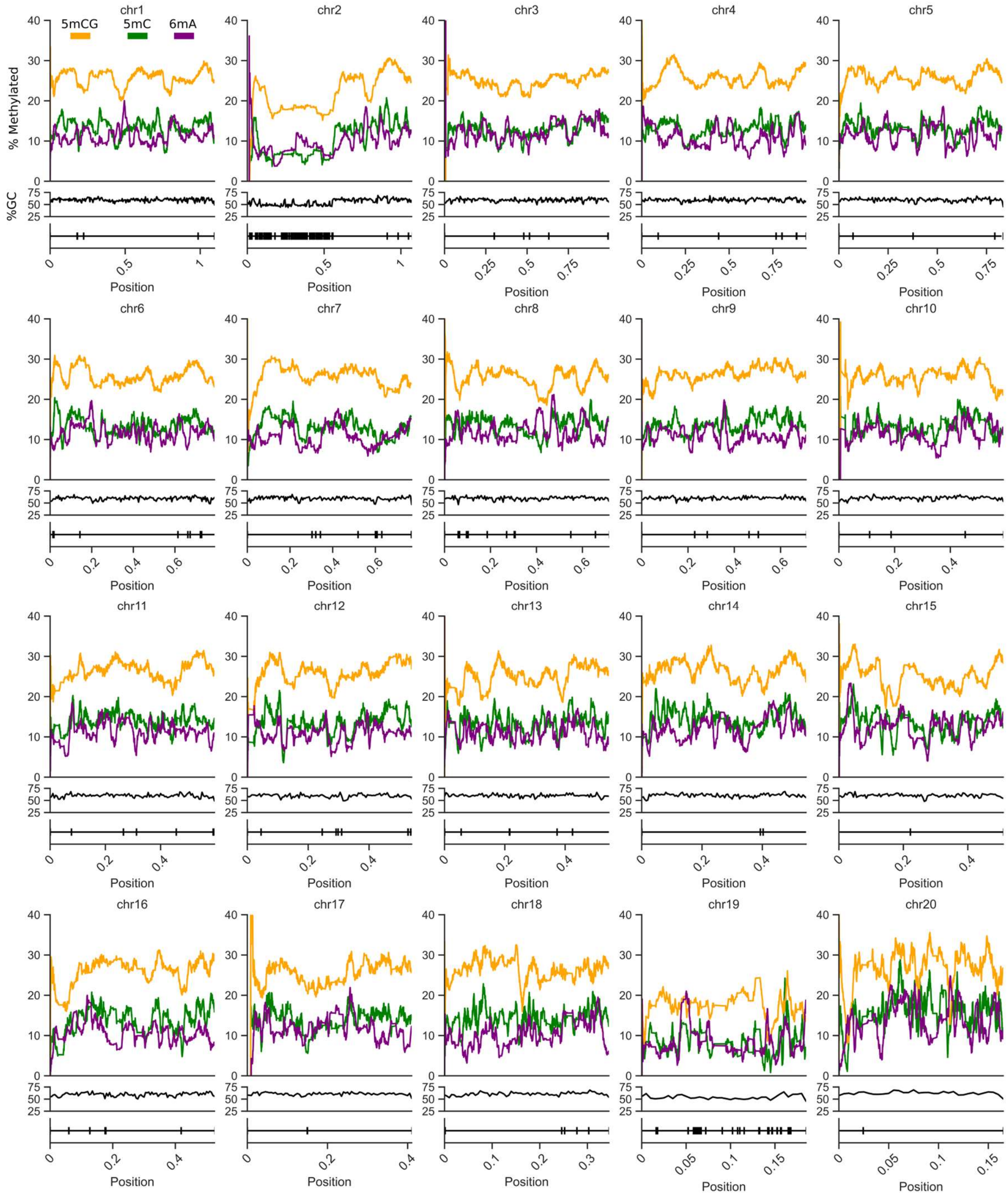


**Figure S7. Hi-C contact maps, transcription, 5mCG methylation, insulation, GC for all chromosomes in *O. tauri*.** Tracks of percentage 5mCG (orange), 5mC (green), 6mA (purple) base modifications over a sliding mean average window of length of chromosome/200 for each nuclear *O. tauri* chromosome. Followed by a track for GC-content sliding mean window of 10kb and a track for transposable element annotation (TEs) across nuclear chromosomes of *O. tauri* RCC4221. Hydroxy-methylation removed as negligible.

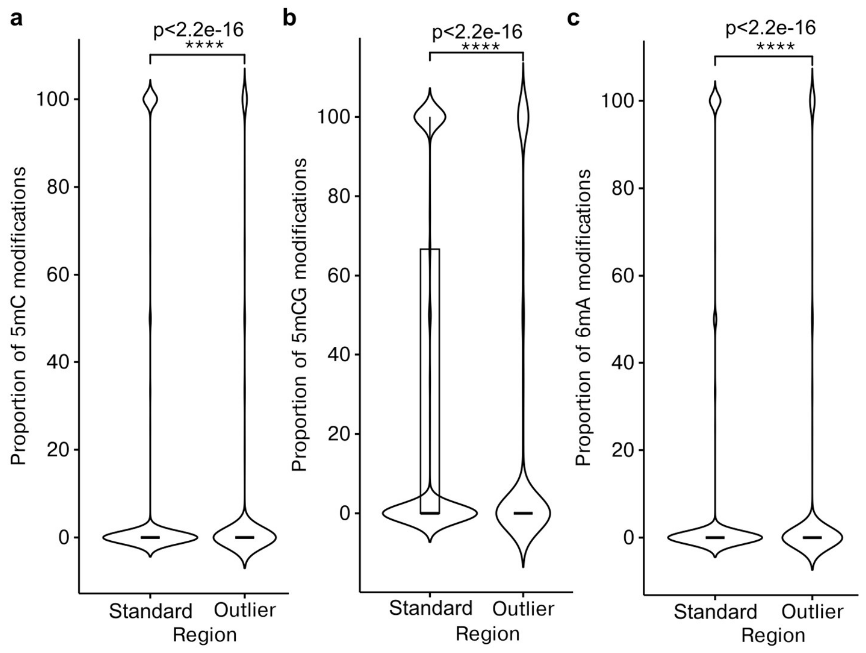


**Figure S8. Decreased methylation across BOC1 and SOC1 of *Ostreococcus tauri* RCC4221.** (**a—c**), Distribution of proportion of cytosine and adenine modifications across low-GC content region of BOC and SOC compared to other nuclear regions. (**a**), 5mC modifications (Wilcoxon rank sum test, p<2.2e-16). (**b**), 5mCG modifications (Wilcoxon rank sum test, p<2.2e-16). (**c**), 6mA modifications (Wilcoxon rank sum test, p<2.2e-16).

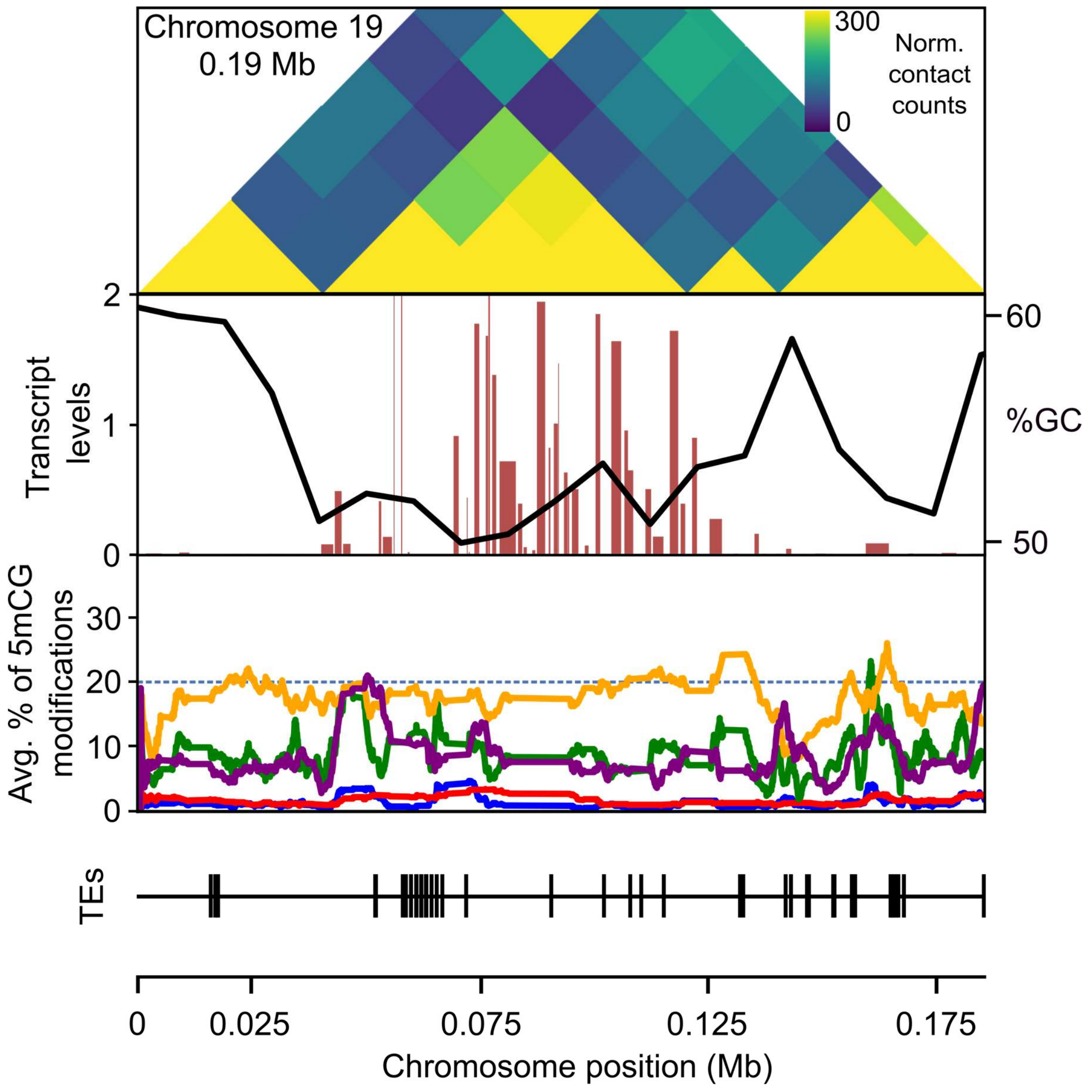


**Figure S9. Elevated transcript abundance and appearance of SOC1 across TAD region on *O. tauri* RCC4221 Chromosome 19.** Hi-C contact map for chromosome 19, binned at a 20 kb resolution. Transcript levels plotted alongside GC in first track. Second track contains a rolling average over a scaled window size of the proportion of 5mCG (orange), 5mC (green), 6mA (purple), 5hmCG (red), 5hmC (blue) modifications. Third track details the transposable elements computationally annotated across the chromosome.

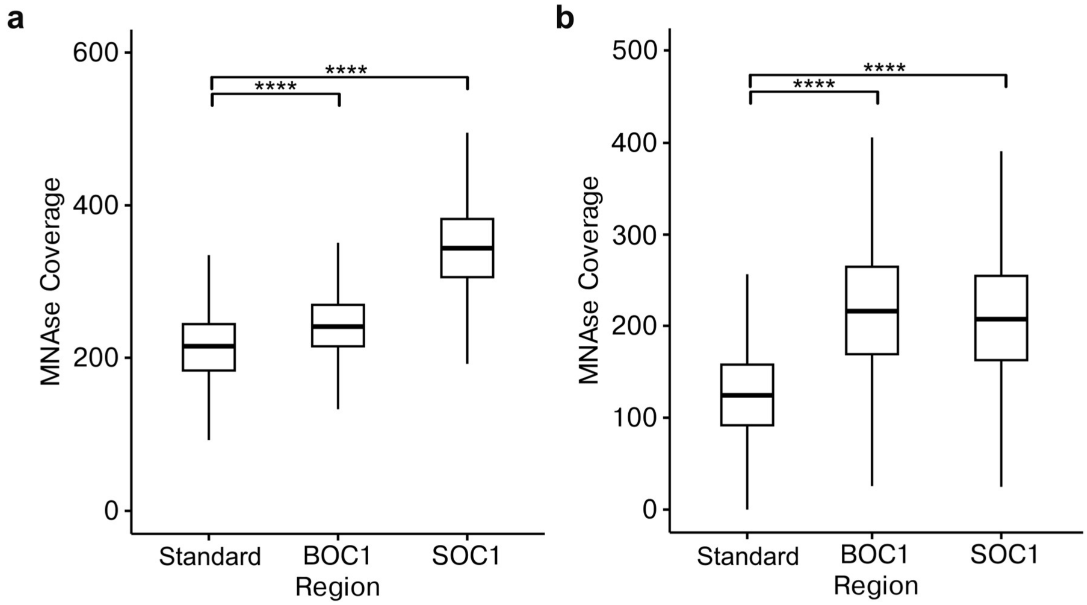


**Figure S10. Increased MNase Coverage across BOC1 and SOC1 of *Ostreococcus lucimarinus* CCE9901 and *Micromonas pusilla* CCMP1545.** (**a**), *O. lucimarinus*. (**b**), *M. pusilla*. (Kruskal-wallis, p<0.001). (**a—b**), Distribution of MNase coverage across low-GC content region of BOC and SOC compared to standard nuclear regions.

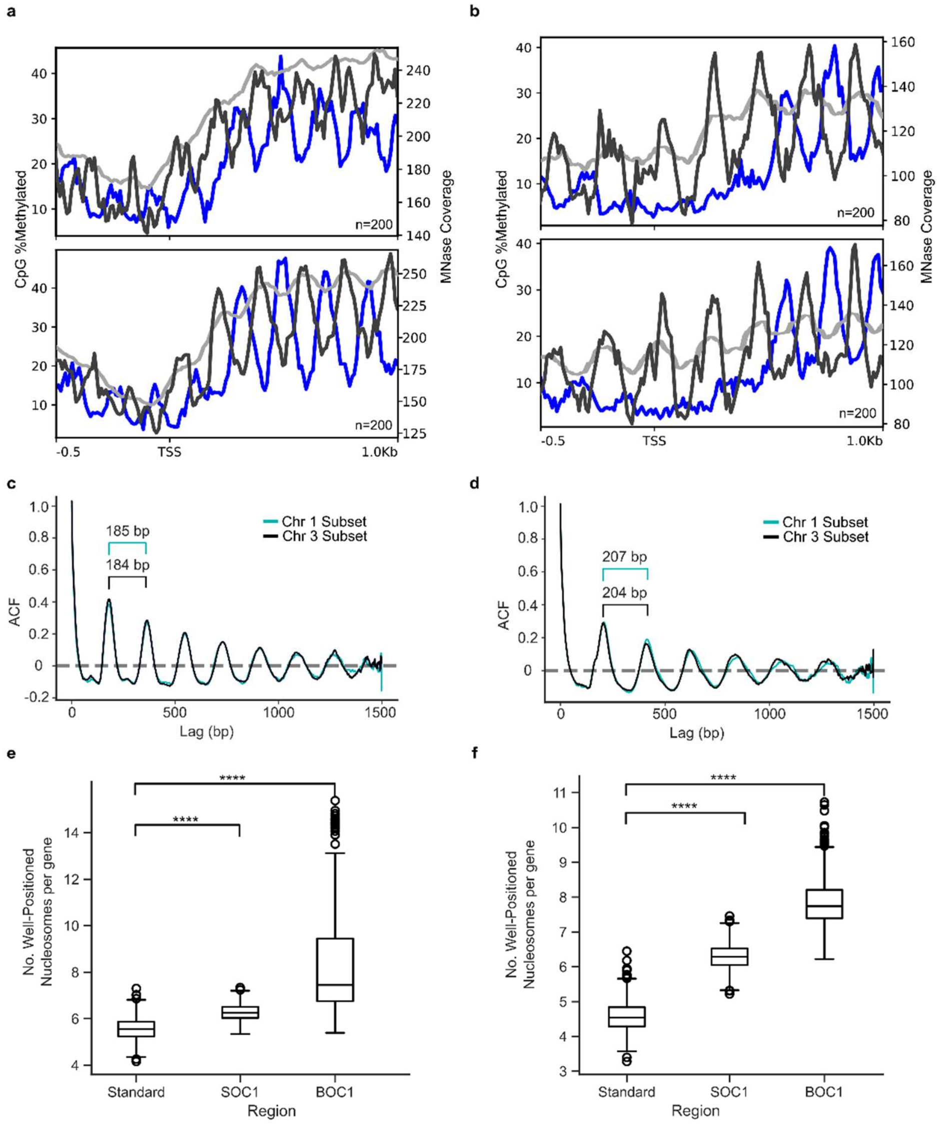


**Figure S11. MNase and CpG across gene bodies on invariant chromosome 1 and chromosome 3.** (**a, c, e**), *Ostreococcus lucimarinus* CCE9901. (**b, d, f**), *Micromonas pusil*la CCMP1545. (**a, b**), Means at each position aligned to transcription start sites are shown for CpG methylation and nucleosomes within a subset of chromosome 1 genes (n=200) above and a subset of chromosome 3 (n=200) genes below. Blue is the percentage of bases which are CpG methylated. Black is nucleosome center counts. Light grey is MNase coverage. (**c, d**), The autocorrelation function estimates of nucleosome center counts are shown for each lag (offset, bp) across a subset of genes on chromosome 1 and 3. Period is annotated in bp. (**e, f**), The distributions of the number of well positioned (non-fuzzy) nucleosomes per genes randomly sampled (n=100, rep=1000) from BOC1, SOC1 and Standard regions ($p<0.001$,Kruskal-Wallis).

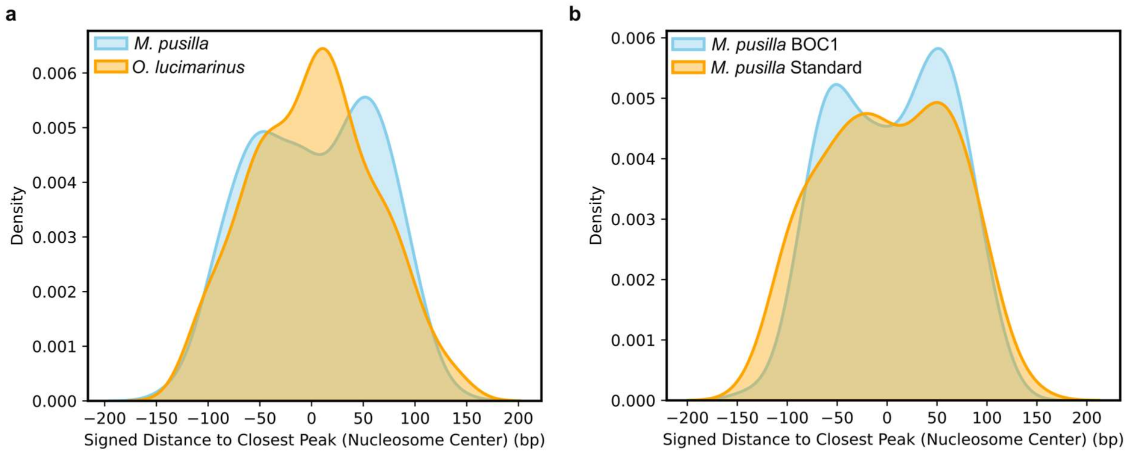


**Figure S12. Occurrences of TEs proximal to nucleosome centers.** (**a**), Kernel density estimation (KDE) plot showing the distribution of transposable elements (TEs) relative to nucleosome centers in *Micromonas pusilla* (blue) and *Ostreococcus lucimarinus* (orange). The x-axis represents the signed distance (in base pairs) from the closest nucleosome center, with negative values indicating upstream positioning and positive values indicating downstream positioning. (**b**), KDE plot comparing *M. pusilla* TEs occurring on BOC1 (blue) versus the standard genome (orange), highlighting differences in their distribution around nucleosome centers. Shaded regions represent density estimates, with peak heights reflecting relative TE frequency near nucleosomes.

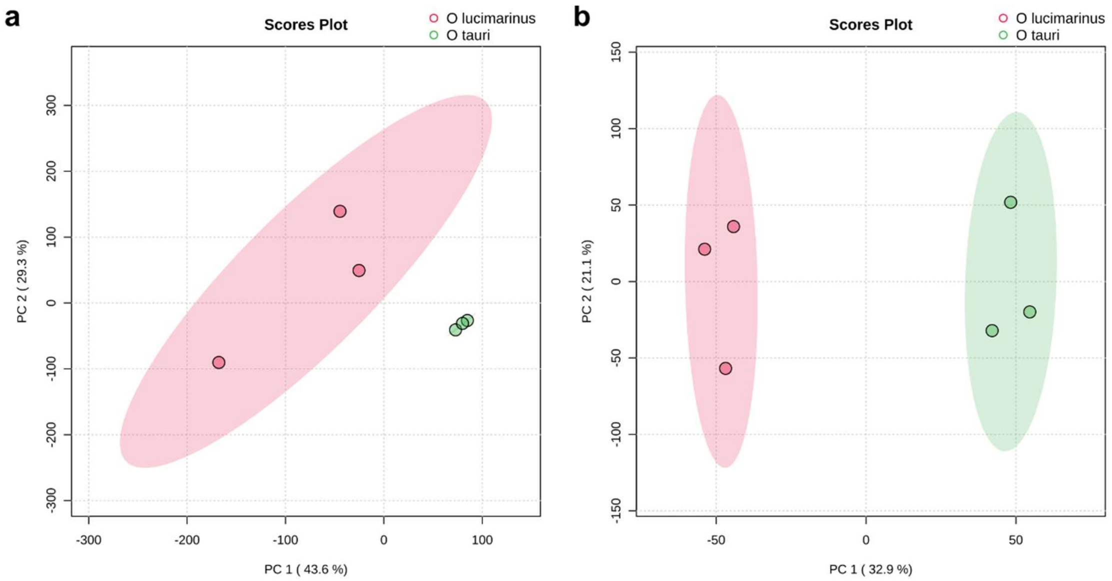


**Figure S13. Metabolome PCA.** (**a,b**), PCA plots describing metabolome diversity across *O. lucimarinus* and *O. tauri*, created using Metaboanalyst. (**a**), Positive ion mode. (**b**), Negative ion mode.

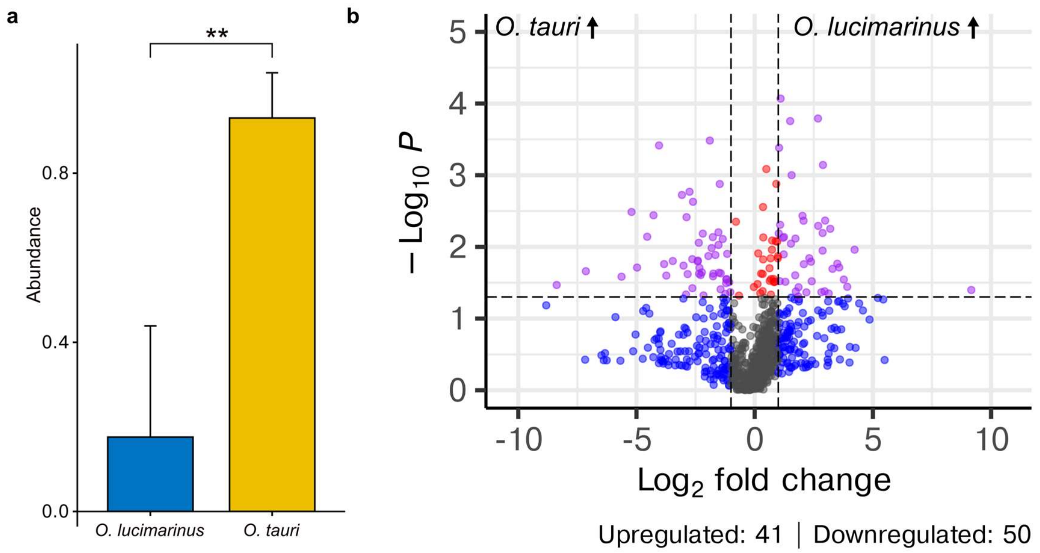


**Figure S14. Putative Polyketide Synthase Product.** (**a**), Bar chart showing differential abundance of putative PKS product ($p<0.005$, one-sided t-test, Alignment ID #3563, Dataset S16–18). (**b**), Volcano plot showing differential metabolite abundances between *O. lucimarinus* (CCE9901) and *O. tauri* (RCC4221) across 895 unidentified negative ion features, revealing 50 features enriched in *O. tauri* versus 41 enriched in *O. lucimarinus*. Fold-change thresholds and statistical significance (ANOVA, $p < 0.05$) are indicated by dashed lines. Gray dots represent non-significantly different metabolites.

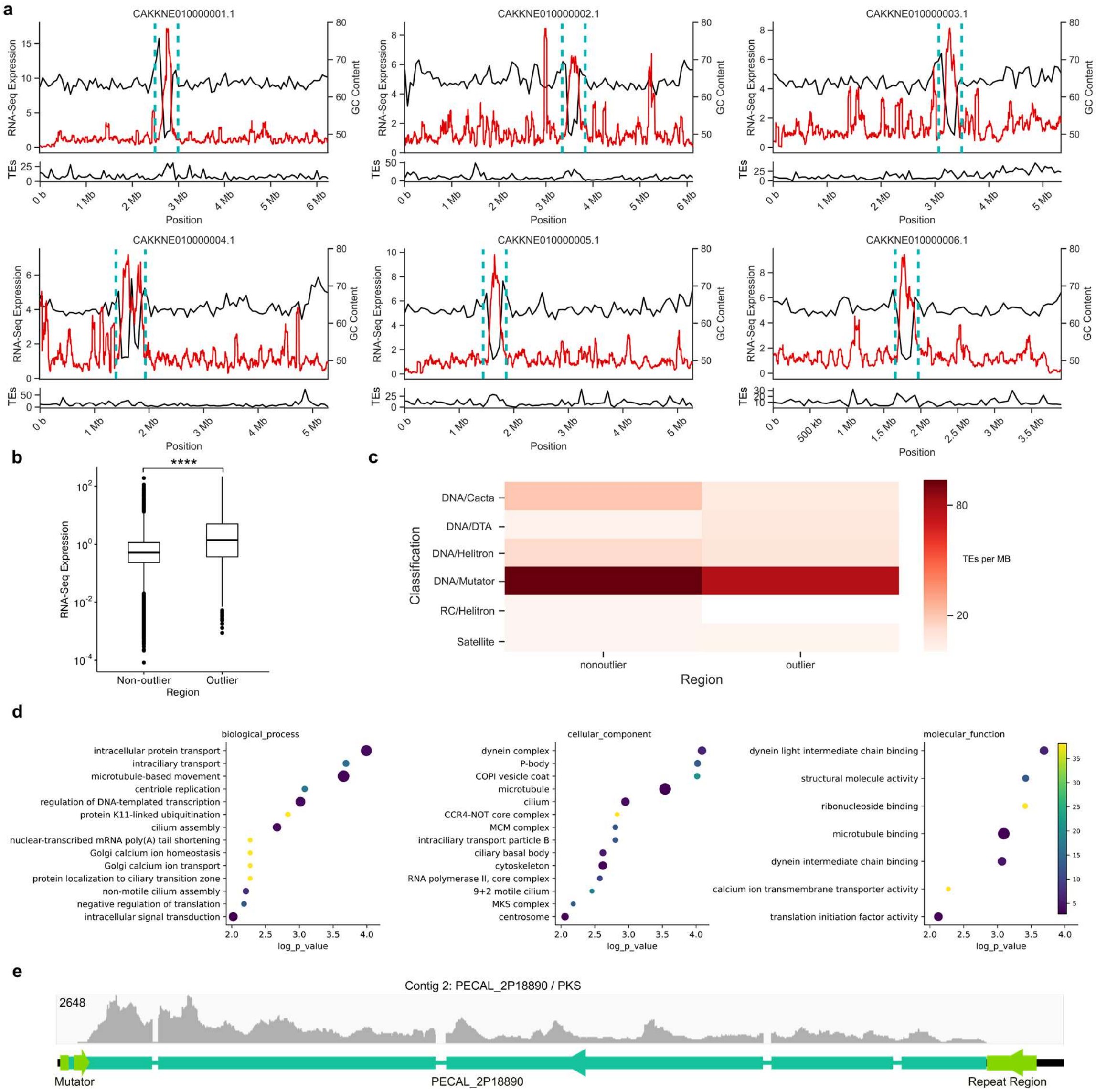


**Figure S15. Pelagomonas calceolata outlier regions.** (**a**), Chromosome 1 – 6. Normalised transcript levels (red) alongside %GC content track (black) - below is a track of the sliding window mean of the number of TEs detected by EDTA. Cyan dotted line represents outlier region. (**b**), Comparative analysis of gene expression in outlier and standard regions- (Kruskal-Wallis, $p<0.001$). (**c**), Heatmap of TEs density of each class in standard genome and outlier (low %GC region). (**d**), Gene ontology terms enriched in proteins across the outlier region. Cutoff p-value ($p<0.01$). Faceted by biological processes, cellular component, molecular function. (**e**), Track of RNA-Seq coverage of PECAL_2P1889 alongside MULEs-transposons.

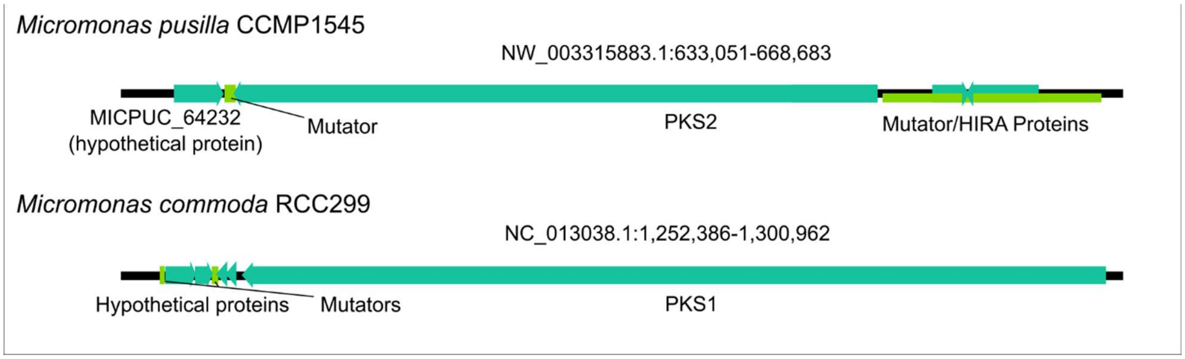


**Figure S16. Mutator transposon co-incidence with giant-type PKS in species other than RCC4221.** Genes coloured in cyan, transposons coloured in green. (**a**), RCC299. (**b**), CCMP1545.

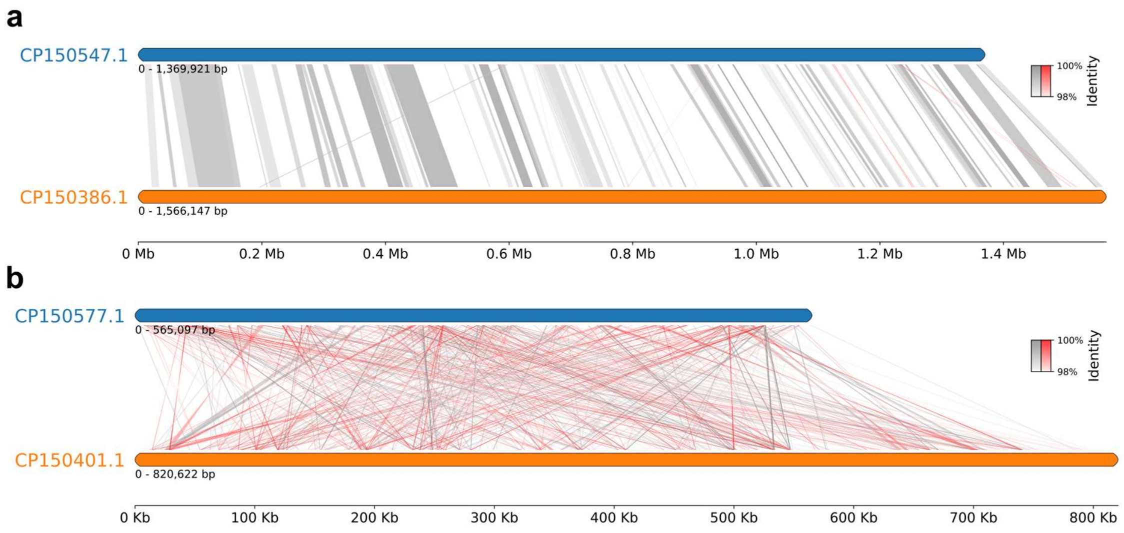


**Figure S17. Homology across invariant and outlier chromosomes of *Pycnococcus provasolii* (top) and *Pseudoscourfieldia marina* (bottom).** (**a**), Homology of an example pair of invariant chromosomes. (**b**), Homology of outlier (low %GC content and high TE occurence) chromosomes.

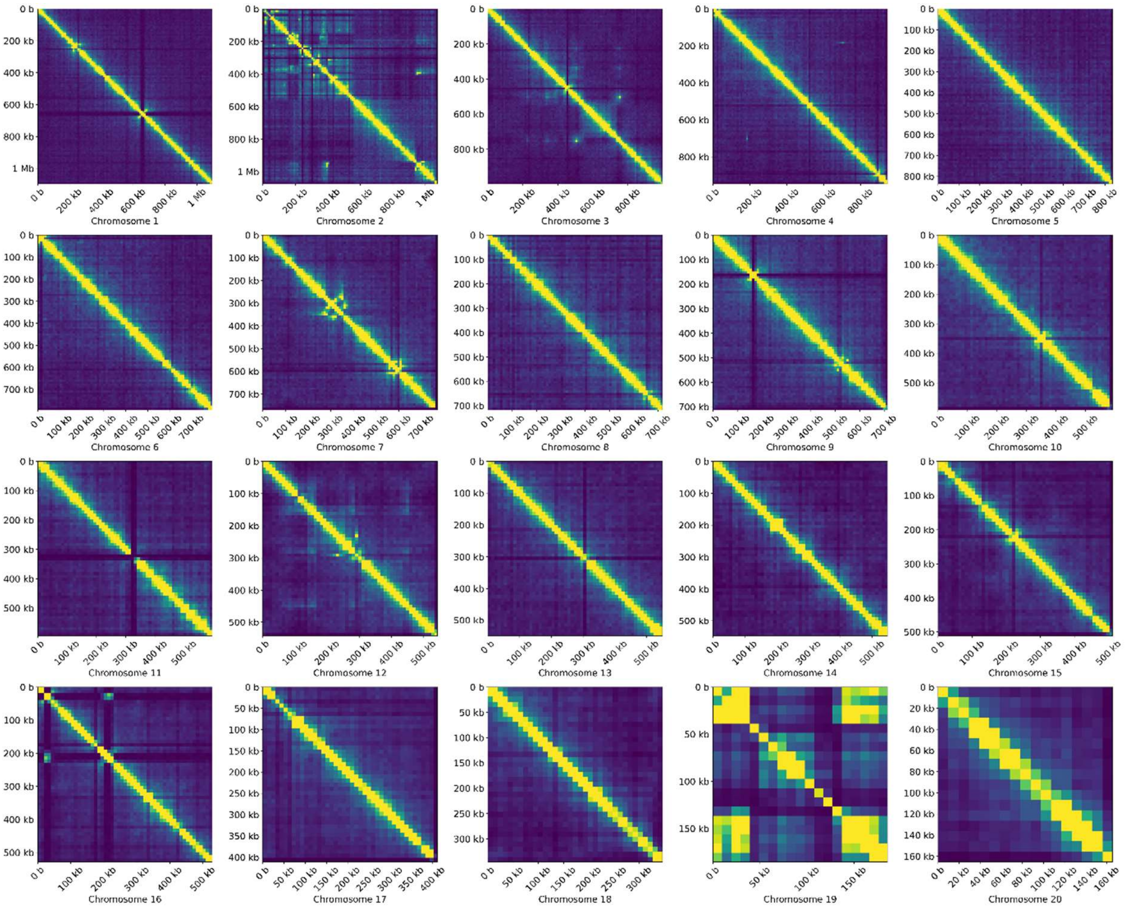


**Figure S18. Hard-masked reference Hi-C contact maps.** Hi-C contact maps made using hard-masked reference genome, 10 kb resolution for the entire *O. tauri* genome.

**a**

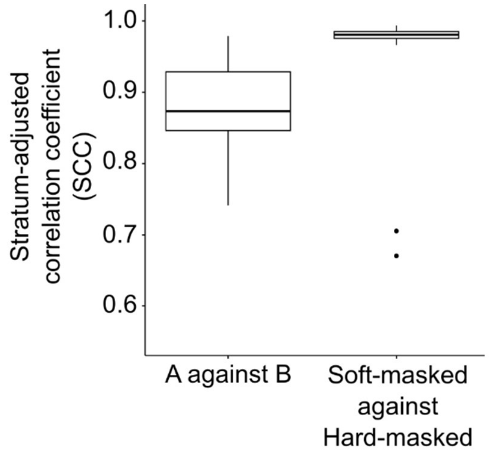


**b**

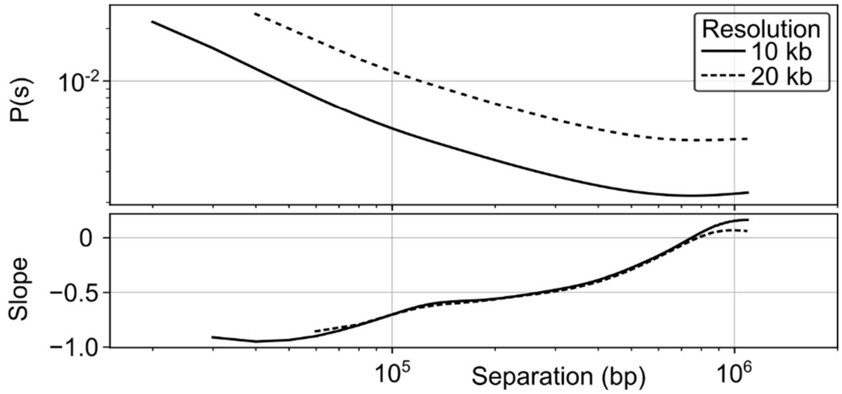


**Figure S19. Contact probability and repeatability.** (**a**), Distribution of stratum-adjusted correlation coefficients of chromosomes between Hi-C contact maps; between replicates A and B and between pooled A and B across a soft- and hard-masked reference. (**b**), Dependence of interaction probability on genomic distance aggregated across the entire chromosome architecture. Alongside its derivative with respect to distance.

### Tables

**Table S1.** Details of the algal strains and publicly available datasets used in this study.

| Species | Strain ID | Origin | Isolation year | Genome | RNA-Seq | Hi-C | ONT Methylation | Bisulfite | MNase | Genome Size (Mb) | Number of Genes | Gene Density (Genes/Mb) |
|---|---|---|---|---|---|---|---|---|---|---|---|---|
| *Ostreococcus tauri* | RCC4221 | Thau Lagoon, FR | 1995 | GCF_000214015.3 | This study | This study | This study | N/A | N/A | 12.9 | 7,833 | 607.2 |
| *Ostreococcus lucimarinus* | CCE99901 / RCC3401 | La Jolla, California, USA | 2013 | GCF_000092065.1 | SRR847308(1) | N/A | N/A | SRR847306 (1) | SRR847307 (1) | 13.2 | 7,640 | 578.8 |
| *Ostreococcus* sp. Clade B | RCC809 | Tropical Atlantic | 1991 | JGI v1.0 (Scaffold) | N/A | N/A | N/A | N/A | N/A | 13.3 | 7,492 | 563.3 |
| *Ostreococcus* sp. Clade C | RCC1115 | Leucate Lagoon, FR | 2006 | GCA_002158475.1 | N/A | N/A | N/A | N/A | N/A | 14.8 | 8,205 | 554.4 |
| *Bathycoccus prasinos* | RCC1105 | Banyuls Bay, FR | 2006 | GCF_002220235.1 | SRR1300453(2) | N/A | N/A | N/A | N/A | 15.0 | 7,967 | 531.1 |
| *Micromonas pusilla* | CCMP1545 / RCC834 | Plymouth, UK | 1950 | JGI v3.2 | (3) | N/A | N/A | SRR847303 (1) | SRR847304 (1) | 22.0 | 10,248 | 465.8 |
| *Micromonas commoda* | RCC299 | Equatorial Pacific, FR | 1998 | GCF_000090985.2 | (4) | N/A | N/A | N/A | N/A | 21.0 | 10,125 | 482.1 |
| *Mantoniella tihuanana* | RCC11003 | South China Sea, CN | 2023 | PRJNA1044521 | N/A | (5) | N/A | N/A | N/A | 39.5 | 11,115 | 281.4 |
| *Pelagomonas calceolata* | RCC100 / RCC697 | North Pacific, Pacific Ocean | 1973 / 2003 | GCA_918797485.1 | ERR3497221 ERR3497222 (6, 7) | (7) | N/A | N/A | N/A | 32.4 | 16,525 | 510.0 |
| *E. pelagica* | UHM3201 | ALOHA, North Pacific Ocean | 2019 | GCA_946965055.2 | N/A | N/A | N/A | N/A | N/A | 60.3 | N/A | N/A |
| *M. coccoides* | CCAP251/1B | Cambridge, UK | 1951 | GCA_963854735.1 | N/A | N/A | N/A | N/A | N/A | 22.3 | N/A | N/A |
| *P. marina* | K-0017 | Oslofjord, Norway | 1983 | GCA_049488355.1 | N/A | N/A | N/A | N/A | N/A | 35.4 | 15,482 | 437.7 |
| *P. provasolii* | CCMP1203 | North Atlantic | 1978 | GCA_049487715.1 | N/A | N/A | N/A | N/A | N/A | 31.5 | 14,349 | 455.5 |

**Table S2**. Low-GC regions on outlier chromosomes (BOC1/SOC1) in Mamiellales (chromosome:start-end).

| *Species/Strain* | *BOC1* | *SOC1* |
|---|---|---|
| *Bathycoccus prasinos* RCC1105 | 14:236,365-624,661 | 19:1-146,238 |
| *Micromonas pusilla* CCMP1545 | 2:60,000-1,733,623* | 19:1-245,704 |
| *Micromonas commoda* RCC299 | 1:263,000-1,817,000 | 17:1- 214,782 |
| *Ostreococcus lucimarinus* CCE9901 | 2:345,000-709,200 | 18:1-149,386 |
| *Ostreococcus tauri* RCC4221 | 2:1-535,000** | 19:55,000-145,000 |
| *Ostreococcus tauri* RCC1115 | 2:150,000-625,000 | 19:1-369,517 |
| *Mantoniella tinhauana* RCC11003 | 2:1,100,000-1,500,000 and 3,000,000 – 3,191,000 | 16:1-310,000 |
| *Ostreococcus* sp. RCC809 | 2: 190,000-505,000 | 18:1-101,961 |

**Table S3**. Correlation between transcript levels and GC content of CDS in *O. tauri* RCC4221*, M. pusilla, M. commoda,* and *B. prasinos* (Spearman's Rank); outlier chromosomes are in **bold.**

| ***chromosome*** | ***p-value*** | ***correlation*** |
|---|---|---|
| | <u>*Ostreococcus tauri* RCC4221</u> | |
| *Chr 1* | 9.193563408752E-23 | -0.366016337734573 |
| ***Chr 2*** | **6.36070819896924E-69** | **-0.66123717210081** |
| *Chr 3* | 6.0557017832092E-11 | -0.260828560220564 |
| *Chr 4* | 2.41217952036462E-24 | -0.41189743117694 |
| *Chr 5* | 1.79552866740855E-09 | -0.261582092461369 |
| *Chr 6* | 2.44394682461919E-09 | -0.271558533584793 |
| *Chr 7* | 5.34058738082654E-10 | -0.276550767710674 |
| *Chr 8* | 1.78261589205752E-11 | -0.316039659786554 |
| *Chr 9* | 9.90218890260923E-11 | -0.295629663567523 |
| *Chr 10* | 2.00787265293745E-12 | -0.360919015773763 |
| *Chr 11* | 3.95199813628221E-15 | -0.411044056868187 |
| *Chr 12* | 7.85332241420064E-10 | -0.335503581607924 |
| *Chr 13* | 5.54123168423855E-12 | -0.375458452239277 |
| *Chr 14* | 1.83465459718988E-11 | -0.351874130368195 |
| *Chr 15* | 4.24845777609905E-14 | -0.409084174287644 |
| *Chr 16* | 2.55333439719958E-11 | -0.369830531679627 |
| *Chr 17* | 1.68814729348831E-05 | -0.263872599808177 |
| *Chr 18* | 4.81543163280944E-09 | -0.394528808280144 |
| ***Chr 19*** | **0.00576848722169023** | **-0.317959481791181** |
| *Chr 20* | 0.000237377090499083 | -0.359691639131982 |
| *Whole Genome* | 7.81552210734339E-242 | -0.366066384736986 |
| | <u>*Micromonas pusilla* CCMP1545</u> | |
| *Scaffold 1* | 4.51559388540416E-61 | -0.477091668432911 |
| ***Scaffold 2*** | **2.58801852972349E-53** | **-0.521738809657552** |
| *Scaffold 3* | 8.34278074397388E-45 | -0.441330208935959 |
| *Scaffold 4* | 1.70177359852189E-32 | -0.408306129128437 |
| *Scaffold 5* | 6.18782887130621E-38 | -0.454198632868908 |
| *Scaffold 6* | 2.666284991579E-27 | -0.426159644002293 |
| *Scaffold 7* | 2.60869365753814E-40 | -0.514111515884707 |
| *Scaffold 8* | 5.08141865672817E-24 | -0.40205175507049 |
| *Scaffold 9* | 5.6306898313881E-33 | -0.489435822113232 |
| *Scaffold 10* | 6.61894117554731E-22 | -0.415020003298428 |
| *Scaffold 11* | 4.95037973143714E-14 | -0.370426040958652 |
| *Scaffold 12* | 6.72344571957175E-30 | -0.505356260644261 |
| *Scaffold 13* | 1.70942233188109E-13 | -0.36034062056912 |
| *Scaffold 14* | 4.6096108508673E-19 | -0.448993750786784 |
| *Scaffold 15* | 1.85444816185892E-15 | -0.409980910517244 |
| *Scaffold 16* | 2.2506691343788E-06 | -0.24728121741501 |
| *Scaffold 17* | 1.29785524507579E-12 | -0.405946551780588 |
| *Scaffold 18* | 2.86141205503479E-08 | -0.374762556017973 |
| ***Scaffold 19*** | **1.84553309705539E-09** | **-0.601289869046519** |
| *Scaffold 20* | 0.104088038661828 | -0.8 |
| *Whole Genome* | 0 | -0.491380286145764 |

| | | |
|---|---|---|
| | <u>*Micromonas commoda* RCC299</u> | |
| ***Chr 01*** | **1.84447779265567E-27** | **-0.36193994892644** |
| *Chr 02* | 1.80806148243263E-16 | -0.260674047245901 |
| *Chr 03* | 3.11047357975182E-10 | -0.207429047434713 |
| *Chr 04* | 4.58762848118535E-12 | -0.242219619321925 |
| *Chr 05* | 2.43568197193201E-07 | -0.18690939568654 |
| *Chr 06* | 9.00135201063736E-10 | -0.225442084071665 |
| *Chr 07* | 7.74251799395708E-08 | -0.198452829861527 |
| *Chr 08* | 1.10357689343177E-11 | -0.279425467726778 |
| *Chr 09* | 2.45435126215848E-09 | -0.230349827323107 |
| *Chr 10* | 0.000174781469433501 | -0.15808657604513 |
| *Chr 11* | 7.22344772220757E-08 | -0.215400408405163 |
| *Chr 12* | 4.85822044898945E-05 | -0.176657680518658 |
| *Chr 13* | 2.12309257075141E-06 | -0.207557665030598 |
| *Chr 14* | 0.000569426344684391 | -0.16824342615038 |
| *Chr 15* | 0.00324820869906 | -0.15324894902115 |
| *Chr 16* | 3.62238604162457E-06 | -0.260420712803006 |
| ***Chr 17*** | **1.21836562682919E-06** | **-0.503570922569601** |
| *Whole Genome* | 3.69823619315502E-192 | -0.285221041029556 |
| | <u>*Bathycoccus prasinos* RCC1105</u> | |
| *Chr 1* | 1.04E-16 | 0.29863 |
| *Chr 2* | 3.84E-20 | 0.357618 |
| *Chr 3* | 1.47E-06 | 0.199046 |
| *Chr 4* | 3.40E-08 | 0.241135 |
| *Chr 5* | 7.28E-12 | 0.294227 |
| *Chr 6* | 8.26E-10 | 0.265054 |
| *Chr 7* | 6.86E-09 | 0.258236 |
| *Chr 8* | 4.62E-06 | 0.201285 |
| *Chr 9* | 4.90E-10 | 0.281628 |
| *Chr 10* | 4.64E-05 | 0.195321 |
| *Chr 11* | 6.88E-06 | 0.228412 |
| *Chr 12* | 1.61E-08 | 0.281895 |
| *Chr 13* | 0.002185 | 0.176243 |
| ***Chr 14*** | **8.49E-13** | **-0.38302** |
| *Chr 15* | 1.34E-10 | 0.374785 |
| *Chr 16* | 7.93E-05 | 0.244112 |
| *Chr 17* | 8.35E-05 | 0.256452 |
| *Chr 18* | 0.000373 | 0.281391 |
| ***Chr 19*** | **0.151531** | **0.17076** |
| *Whole Genome* | 6.21E-88 | 0.222575 |

**Table S4**. Table of Outlier region information, GC% and TEs/Mb across other minimal algae used in the study.

| *Species* | *Outlier Chr* | *Outlier Region* | *Standard GC%* | *Outlier GC%* | *Standard TEs/Mb* | *Outlier TEs/Mb* |
|---|---|---|---|---|---|---|
| *Epithemia pelagica* | OX337233.1 | 1.6-1.9 Mb | 48.15 | 46.46 | 373.18 | 416.67 |
| *Epithemia pelagica* | OX337238.1 | 2.75-3.1 Mb | 48.15 | 47.4 | 373.18 | 1548.57 |
| *Epithemia pelagica* | OX337240.1 | 2-2.2 Mb | 48.15 | 46.29 | 373.18 | 745 |
| *Marvania coccoides* | OY978099.1 | 1.9-2.312 Mb | 54.97 | 48.55 | 84.62 | 864.08 |
| *Pelagomonas calceolata* | CAKKNE010000001.1 | 2.5-3 Mb | 63.93 | 61.61 | 167.91 | 266 |
| *Pelagomonas calceolata* | CAKKNE010000002.1 | 3.4-3.75 Mb | 63.93 | 57.21 | 167.91 | 377.14 |
| *Pelagomonas calceolata* | CAKKNE010000003.1 | 3.1-3.5 Mb | 63.93 | 59.05 | 167.91 | 290 |
| *Pelagomonas calceolata* | CAKKNE010000004.1 | 1.4-1.9 Mb | 63.93 | 58.47 | 167.91 | 268 |
| *Pelagomonas calceolata* | CAKKNE010000005.1 | 1.4-1.9 Mb | 63.93 | 60.44 | 167.91 | 260 |
| *Pelagomonas calceolata* | CAKKNE010000006.1 | 1.6-2 Mb | 63.93 | 59.25 | 167.91 | 260 |
| *Pseudoscourfieldia marina* | CP150401.1 | Whole Chr (0.82 Mb) | 57.05 | 49.77 | 283.48 | 1208.84 |
| *Pycnococcus provasolii* | CP150577.1 | Whole Chr (0.57 Mb) | 57.51 | 49.2 | 18.77 | 424.71 |

**Supplementary Information**

**Dataset S1 – Table of Gene Ontology Enrichment on BOC1 Region.**

**Dataset S2 – Table of Gene Ontology Enrichment on SOC1 Region.**

**Dataset S3 – Table of Transposable Element class of Mamiellales genome across outlier and standard regions.**

**Dataset S4 – Table of genomic statistics of TE, Intron and Transcript abundance in Mamiellales genomes.**

**Dataset S5 – Table of annotated TE elements across RCC4221 Reference Genome.**

**Dataset S6 – Table of Transcript abundance across standard and outlier regions in Mamiellales genomes.**

**Dataset S7 – Table of Splice Junctions across chromosomes in Mamiellales.**

**Dataset S8 – Table of Hi-C Statistics for *O. tauri* and *M. tinhauana*.**

**Dataset S9 – Table summarizing chromatin loops detected across *O. tauri* RCC4221.**

**Dataset S10 – Table of interchromosomal interactions between standard and outlier chromosomes.**

**Dataset S11 – Table of ONT methylation data, % modified, for *O. tauri*.**

**Dataset S12 – Table of CpG and MNAse data for *O. lucimarinus* and *M. pusilla*.**

**Dataset S13 – Table of nucleosome_dynamics summary for *O. lucimarinus* and *M. pusilla* using BOC1, and subsets of chromosomes 1 and 3.**

**Dataset S14 – Table of PKS architectures across genomes used in article.**

**Dataset S15 – Table of ANOVA summary stats for detected metabolites in *O. tauri* and *O. lucimarinus*.**

**Dataset S16 – Table of detected metabolites in positive ion mode in *O. tauri* and *O. lucimarinus*.**

**Dataset S17 – Table of detected metabolites in negative ion mode in *O. tauri* and *O. lucimarinus*.**

**Dataset S18 – Table of further Sirius analysis of differential unknown metabolites.**

**Dataset S19 – Table of EDTA results for Pseudoscourfieldia marina, Pycnococcus provasolii and Pelagomonas calceolata.**
